# Widespread chromatin context-dependencies of DNA double-strand break repair proteins

**DOI:** 10.1101/2022.10.07.511243

**Authors:** Xabier Vergara, Anna G. Manjón, Ben Morris, Ruben Schep, Christ Leemans, Mathijs A. Sanders, Roderick L. Beijersbergen, René H. Medema, Bas van Steensel

**Affiliations:** Division of Gene Regulation, Netherlands Cancer Institute, 1066 CX Amsterdam, The Netherlands; Division of Cell Biology, Netherlands Cancer Institute, 1066 CX Amsterdam, The Netherlands; Division of Molecular Carcinogenesis, Netherlands Cancer Institute, 1066 CX, Amsterdam, The Netherlands; Oncode Institute, Netherlands Cancer Institute, 1066 CX, Amsterdam, The Netherlands; Department of Hematology, Erasmus MC Cancer Institute, Rotterdam, The Netherlands; Cancer, Ageing and Somatic Mutation (CASM), Wellcome Sanger Institute, Hinxton, UK

## Abstract

DNA double-strand breaks are repaired by multiple pathways, including non-homologous end-joining (NHEJ) and microhomology-mediated end-joining (MMEJ). The balance of these pathways is dependent on the local chromatin context, but the underlying mechanisms are poorly understood. By combining knockout screening with a dual MMEJ:NHEJ reporter inserted in 19 different chromatin environments, we identified dozens of DNA repair proteins that modulate pathway balance dependent on the local chromatin state. Proteins that favor NHEJ mostly synergize with euchromatin, while proteins that favor MMEJ generally synergize with distinct types of heterochromatin. BRCA2 is an example of the former, which is corroborated by chromatin-dependent shifts in mutation patterns of BRCA2^-/-^ cancer genomes. These results uncover a complex network of proteins that regulate MMEJ:NHEJ balance in a chromatin context-dependent manner.

**ONE SENTENCE SUMMARY:** A multiplexed screen reveals how dozens of proteins sense the local chromatin context to tune the balance between two DNA repair pathways.

## MAIN TEXT

### Background: chromatin context effects on DSB repair pathways

DNA double-strand breaks (DSB) are repaired by multiple repair pathways such as non-homologous end-joining (NHEJ), homologous recombination (HR) and microhomology-mediated end joining (MMEJ). These pathways act in an equilibrium that is referred to as the DNA repair pathway balance(reviewed in *1*). Defects in this balance can compromise genome stability, but also offer opportunities for therapy, particularly in cancer (*2*). Pathway balance is influenced by several factors, including cell cycle (*3*), break complexity (*4*) and the chromatin context in which a DSB occurs (*5, 6*). The latter is generally attributed to molecular interactions between specific repair proteins and distinct chromatin proteins, in some instances regulated by posttranslational modifications (*7-9*). Such local interactions can alter the recruitment of the repair protein to a DSB, or modulate its activity in the repair process. Yet, the overall extent and the principles of this interplay between chromatin and repair proteins have remained poorly studied. Here, by screening hundreds of DNA repair proteins, we uncover that chromatin context has a widespread influence on the relative contribution of specific DNA repair proteins to repair pathway balance.

### Experimental design

We focused on the balance between NHEJ and MMEJ, which are two of the main mutagenic DSB repair pathways, particularly for DSBs generated during CRISPR editing (*10*). We applied a sequencing-based assay that determines the MMEJ:NHEJ balance after induction of a DSB by Cas9, with high accuracy and in multiple genomic loci in parallel(*6*). For this we employed a human K562 cell line with 19 barcoded Integrated Pathway Reporters (IPRs) inserted throughout the genome (**Fig. 1A**). Importantly, the integration sites represent all major known chromatin types (*6*) (see below). In this cell line we conducted three biological replicates of a 96-well CRISPR/Cas9 screen to knock out (KO) 519 proteins that had previously been linked to at least one DNA repair pathway (**Fig. 1A & Fig. S1A-C; Table S1-S2; Detailed protocol in methods**). For each KO we then induced a DSB in all IPRs; after 72 hours to allow repair to occur, we isolated genomic DNA and sequenced the IPRs to determine the MMEJ:NHEJ balance as the ratio between the signature indels +1 (NHEJ_ins_) and -7 (MMEJ_del_) (**Table S3**) (*6*). For each IPR−KO combination we then computed the log_2_ fold change in MMEJ:NHEJ balance [Δlog_2_MMEJ:NHEJ] relative to the average of a set of 33 mock KO control samples (in which gRNA was omitted in the KO step). We averaged the results of three replicates, resulting in a 519 × 19 matrix of Δlog_2_MMEJ:NHEJ scores (**Table S4**). These scores reflect the contribution of each tested protein to the MMEJ:NHEJ balance in 19 well-characterized chromatin contexts (**Fig. S1D**).

**Fig. 1.**
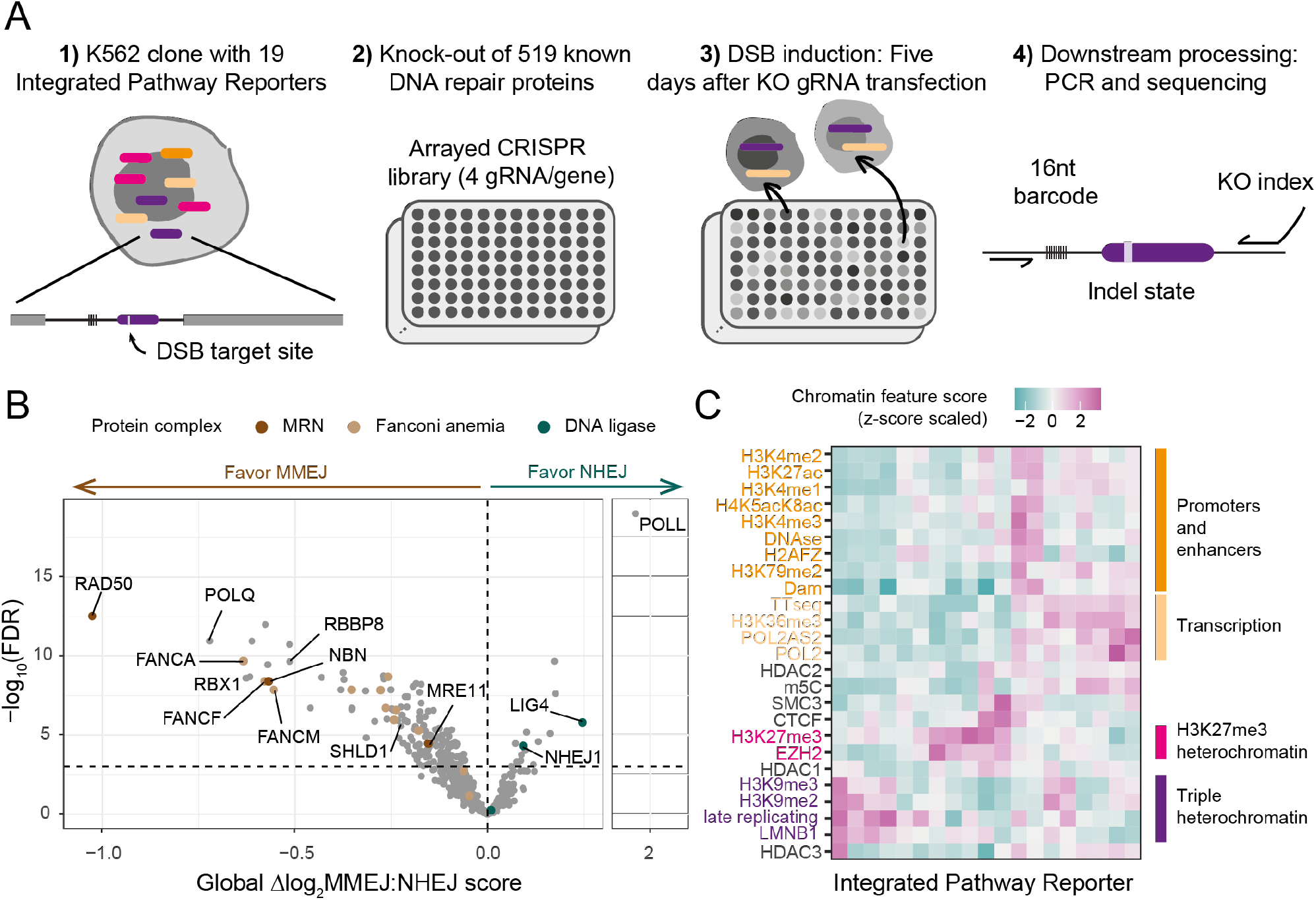
Multiplexed CRISPR screen to assess chromatin context-dependencies of DNA repair proteins. (**A**) Overview of screen design. See main text for explanation. (**B**) Volcano-plot of global Δlog_2_MMEJ:NHEJ scores (mean value of 19 IPRs). Horizontal dotted line shows significance threshold (FDR = 0.001). Labels mark proteins highlighted in the text. (**C**) Heatmap of levels of 25 chromatin features at the 19 IPR sites. Four major chromatin states and their defining features are highlighted in distinct colors.

### Repair proteins affecting pathway balance globally

We first assessed the impact of the tested proteins on the global MMEJ:NHEJ balance, i.e., irrespective of the local chromatin context, by evaluating the mean Δlog_2_MMEJ:NHEJ scores of the 19 IPRs. At an estimated false discovery rate (FDR) of 0.001, 149 proteins *favored MMEJ* (**Fig. 1B**), i.e., these proteins either are required for full MMEJ activity or they inhibit NHEJ when present. Among these are known key components of the MMEJ pathway, such as POLΘ (POLQ), proteins of the MRN complex and CtIP (RBBP8). We also found that several Fanconi anemia (FA) proteins (e.g., FANCA, FANCF, FANCM, FANCD2), which are central proteins of inter-strand crosslink (ICL) repair, favored MMEJ. Unexpectedly, proteins that either directly (SHLD1) (*11*)) or indirectly (RBX1 (*12*)) limit long-range resection, a key step for HR, favored MMEJ. This suggests that limitation of long-range resection favors MMEJ over NHEJ. Conversely, 16 proteins favored NHEJ globally, including known components of the NHEJ pathway, such as Ligase IV (LIG4) (*13, 14*), XLF (NHEJ1) (*15, 16*) and DNA polymerase lambda (POLL) (*17*). Thus, the screen confirmed several known key proteins in the repair of DSBs generated by Cas9 and other nucleases (*17, 18*).

### Many repair proteins show significant chromatin context-dependency

Next, we asked which proteins exhibited chromatin context-dependency (CCD) of their Δlog_2_MMEJ:NHEJ scores across the 19 IPRs. As we and others previously demonstrated, integrated reporters generally adopt the local chromatin state (*6, 19-21*). We therefore used a set of high-quality epigenome maps from K562 cells (**Table S5**) to infer the levels of 25 chromatin features on each of the IPRs (**Fig. 1C, Table S6**). We then applied a three-step linear modeling approach (see Methods, **Fig. S2-3**) to identify proteins for which the Δlog_2_MMEJ:NHEJ scores correlated significantly with one or more chromatin features. According to this analysis, 89 (17.1%) of all tested proteins showed a significant CCD at 5% FDR cutoff. Of 33 mock KO samples only one (3%) passed this cutoff, confirming the low rate of false positives. These results indicate that a surprisingly large proportion of DNA repair proteins modulate the MMEJ:NHEJ balance with a significant CCD (**Table S7**).

### Distinct patterns of synergies

Next, for each of the identified proteins we asked which chromatin features explain the CCD. For this we considered the slope of linear fits that correlate Δlog_2_MMEJ:NHEJ scores with each individual chromatin feature (see Methods, **Fig. S4**). A synergy score (slope) is positive when the repair protein favors NHEJ with increasing levels of the chromatin feature (**I in Fig. 2A**). We will refer to this as “N-synergy”. When the synergy score is negative, the protein favors MMEJ with increasing levels of the chromatin feature (**II in Fig. 2A**); this we will refer to as “M-synergy”. For example, we found that RAD50 is M-synergistic with Lamin B1 (LMNB1) (**Fig. 2B & Fig. S4D**), indicating that RAD50 favors MMEJ preferentially in regions that interact with the nuclear lamina. A synergy score near zero points to a lack of detectable synergy of the tested pair **(III in Fig 2A**), as exemplified by the repair protein MDC1 and the chromatin feature H2AFZ (**Fig. 2C**).

**Fig. 2.**
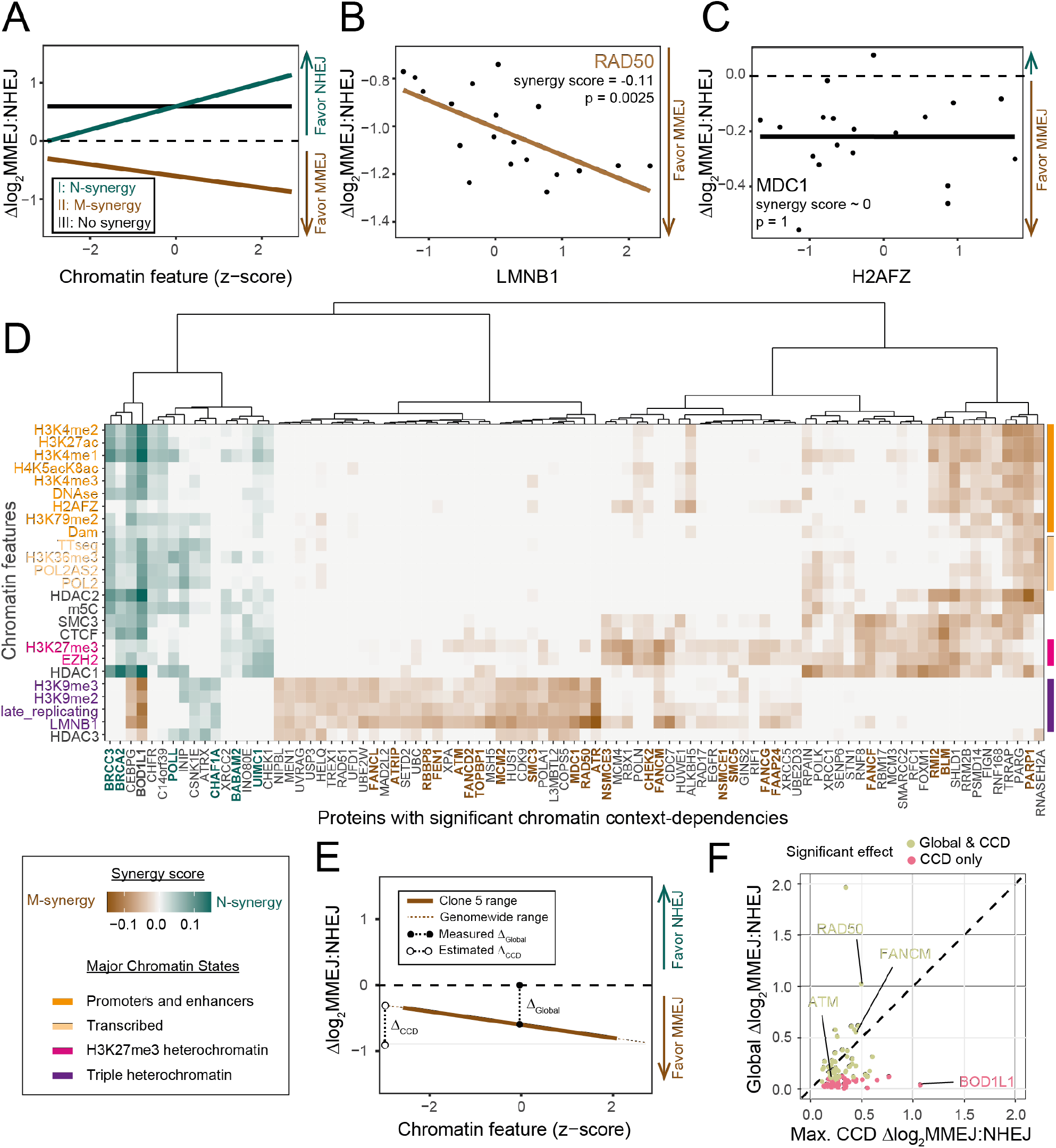
CCDs of 89 DNA repair proteins. (**A**) Illustration of M- and N-synergy concepts. For chromatin feature − protein combinations with N-synergy (I in green) Δlog_2_MMEJ:NHEJ scores increase as the chromatin feature levels increase. For combinations with M-synergy (II in brown) Δlog_2_MMEJ:NHEJ scores decrease as the chromatin feature levels increase. For combinations with no synergy (III in black) Δlog_2_MMEJ:NHEJ scores do not correlate with the individual chromatin features. (**B**) M-synergy example: linear fit (brown line) of RAD50 Δlog_2_MMEJ:NHEJ scores with LMNB1 interaction levels. (**C**) No synergy example: linear fit (black line) of MDC1 Δlog_2_MMEJ:NHEJ scores with H2AFZ levels. (**D**) Heatmap of synergy scores of all 89 proteins with significant CCDs. Proteins (columns) mentioned in the text are highlighted in bold. Chromatin features (rows) are colored and ordered as in **Fig. 1C**. (**E**) Comparison of CCD effect sizes to global effect sizes of the proteins in (D). Global effect sizes are absolute values of global Δlog_2_MMEJ:NHEJ scores as calculated in **Fig. 1B**. CCD effect sizes are the predicted genome-wide dynamic range of Δlog_2_MMEJ:NHEJ values for each protein KO, as a function of variation in chromatin context (see Methods).

### M- and N-synergies: distinct distributions across chromatin types

Hierarchical clustering of the synergy scores of all 89 proteins with significant CCDs revealed striking patterns (**Fig. 2D**). First, 16 proteins have N-synergies while 75 have M-synergies, with two proteins overlapping due to mixed synergies. Thus, proteins with M-synergies are much more prevalent than proteins with N-synergies. This may reflect a higher complexity of the MMEJ pathway compared to the NHEJ pathway (*17, 18, 22*). Second, N-synergies predominantly involve euchromatic features, such as marks of active promoters and enhancers (e.g., H3K4me3 and H3K27ac) and transcription activity (e.g., TT-seq, POL2 and H3K36me3) (**Fig. 2D**). Only a few proteins show N-synergy with heterochromatin, either alone or in combination with a subset of euchromatic marks. Third, M-synergies are divided over three main clusters, with prominent roles for distinct classes of heterochromatin. One cluster of 33 proteins has consistent M-synergy with heterochromatin that is marked by a combination of H3K9me2/3, late replication and interactions with LMNB1. We will refer to this type of heterochromatin as “triple heterochromatin”. A second cluster of 31 proteins is primarily M-synergistic with H3K27me3-marked heterochromatin, often combined with LMNB1; and a third cluster of 11 proteins shows M-synergy with various euchromatin marks, frequently combined with H3K27me3. Thus, the vast majority of M-synergies involve either triple or H3K27me3 heterochromatin, unlike most N-synergies. (**Fig. 2D**). The skewed distribution of M- and N-synergies between heterochromatin and euchromatin provides an explanation for the earlier observation that the MMEJ:NHEJ ratio tends to be higher in heterochromatin (*6*). Interestingly, two proteins (BOD1L1 and CEBPG) are both M- and N-synergistic, indicating that they have opposite roles dependent on the chromatin context (**Fig 2D**). **CCD effects compared to global effects**. Of the 89 proteins with significant CCDs, 46 modulate MMEJ:NHEJ balance globally with preferential impact on specific chromatin contexts (e.g. RAD50, FANCM or ATM). In these cases, CCD and global effects tend to have similar effect sizes (**Fig. 2E-F & Fig. S5**, see Methods). Additionally, 43 proteins only modulate MMEJ:NHEJ balance in specific chromatin contexts (**Fig. 2E-F**). Thus, the magnitude of CCD effects is often similar or larger than the chromatin-independent contributions of individual proteins.

### Interpretation of M- and N-synergies

We note that M-synergy does not necessarily imply that the protein locally boosts MMEJ; it may also locally suppress NHEJ and thereby shift the balance. Similarly, N-synergy may be either due to local activation of NHEJ or local suppression of MMEJ. Furthermore, we emphasize that the synergies as defined here do not necessarily imply a direct molecular link between the repair protein and the chromatin feature; the feature may also be a proxy for an unknown chromatin feature that is closely linked. For this reason, most of our analyses below focus on the major known chromatin states that are represented by one or more features in our dataset. We also note that some hits in our screen can be explained by indirect effects. For example, FOXM1 and EGFR are known to be regulators of various genes that encode DNA repair proteins (*23, 24*), while there is no evidence that they directly mediate DNA repair. Below we highlight findings that are more likely to involve close interactions with chromatin.

### M-synergies of canonical MMEJ proteins

Among canonical components of the MMEJ pathway, several exhibit M-synergy. This includes RAD50 (**Fig. 2B**), CtIP/RBBP8 and FEN1, which show exclusive M-synergy with triple heterochromatin; and PARP1 which has selective M-synergy with euchromatin and H3K27me3 (**Fig. 2D**).

### Proteins that interact tend to have similar CCD patterns

Some proteins that are part of the same complex, such as BLM and RMI2 (*25*), show highly similar M-synergy (**Fig. 2D**). We asked whether this is a general trend among pairs of proteins that are known to physically interact *in vivo* according to the BioGRID database (*26*). We identified n = 118 interacting pairs among proteins with significant CCDs (**Table S8**). Similarities in CCD patterns were significantly higher between physically interacting proteins than expected by random sampling (**Fig. 3A-B**, empirical test p < 0.001). Among the 118 interacting pairs, we even found three ‘cliques’ of at least four proteins that are connected by pairwise physical interactions (**Fig. 3C**). One of these cliques encompasses ATM and its phosphorylation targets MDC1, TOPBP1, and FANCD2. All these proteins show highly similar M-synergies with triple heterochromatin (**Fig. 3D**). TOPBP1 interacting proteins ATRIP and ATR also show M-synergies with triple heterochromatin. In line with this, ATM, TOPBP1, ATR and ATRIP have been previously linked to repair of heterochromatin DSBs (*27, 28*). ***Role of ATM signaling in heterochromatin***. To further investigate the CCD of ATM in heterochromatin, we treated cells with the ATM kinase activity inhibitor KU55933 (**Fig. S6A-B**). ATM inhibition exhibited significant M-synergies with H3K27me3 and interactions with LMNB1, but did not exhibit M-synergies with other triple heterochromatin features. This CCD pattern is more similar to CHEK2, ATM’s main signal transducer (*29*), than ATM itself (**Fig. 3G&I**). This suggests that loss of ATM downstream signaling impacts CCDs differently than losing ATM itself, in line with earlier observations that loss and inhibition of ATM can have different effects (*30, 31*). These data underscore the importance of the ATM signaling axis in repair of DSB in heterochromatin.

**Fig. 3:**
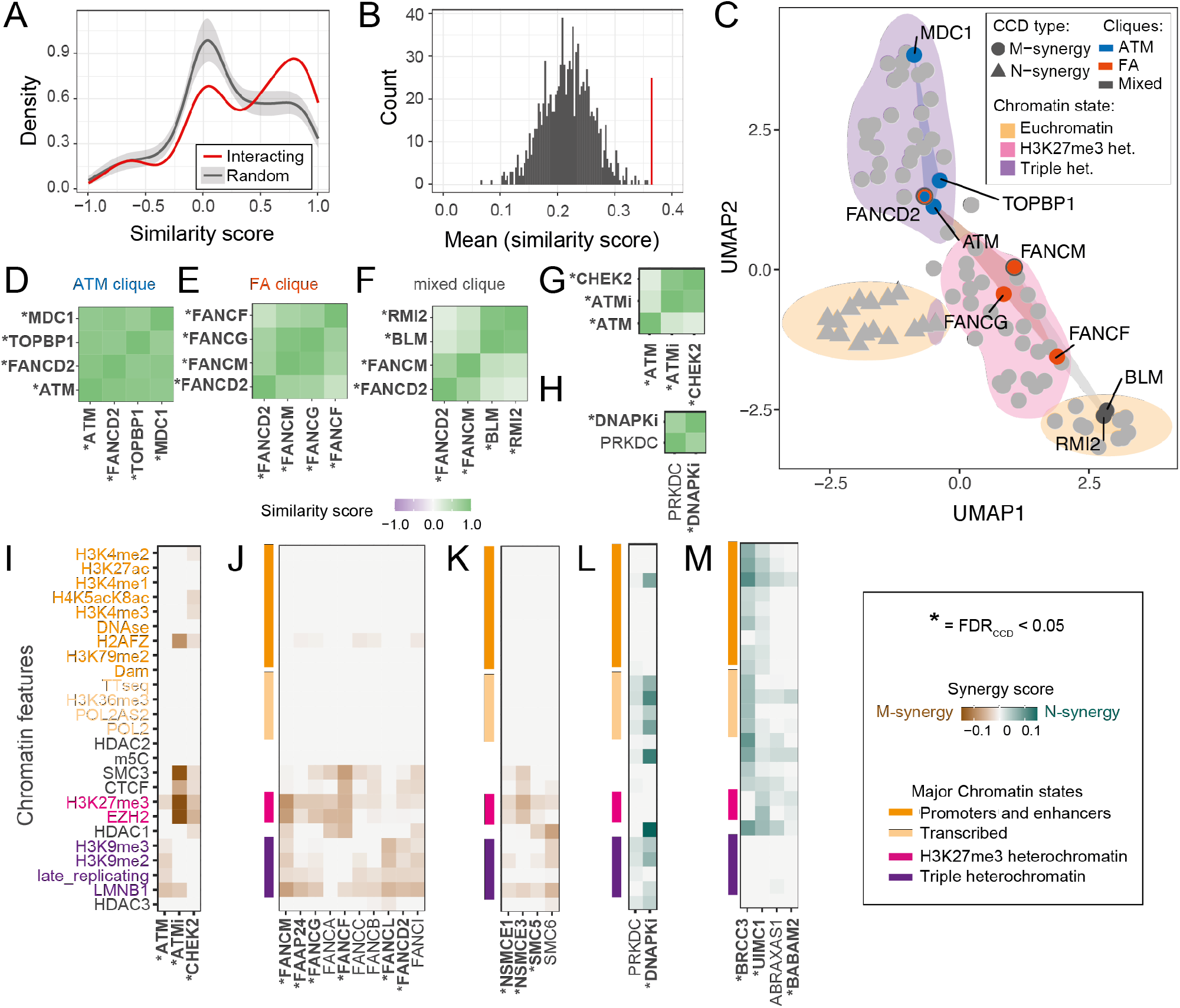
Proteins that physically interact tend to have similar CCD patterns. (**A**) Distributions of pairwise similarity scores for CCD patterns across the 25 chromatin features, between interacting proteins (red; 118 pairs) and between randomly picked protein pairs (mean ± s.d. of 1,000 draws of 118 random pairs). (**B**) Mean similarity score of 118 interacting protein pairs (red line) compared to the distribution of mean similarity scores of 1,000 random draws as in (A) (grey histogram), indicating that high similarities of CCD patterns of interacting protein pairs cannot be explained by random chance (p<0.001). (**C**) Uniform Manifold Approximation and Projection (UMAP) visualization of proteins with CCDs. Each dot represents a protein, with the shape indicating the type of synergy. Color clouds show the major chromatin state that explains each CCD. Three ‘cliques’ of four interacting proteins are shown as colored quadrangles. Proteins shared between multiple cliques are marked by concentric circles with the color of each clique they are part of. (**D-H**) CCD similarity score matrix of proteins in ATM clique (**D**), FA clique (**E**), mixed clique (**F**), ATM signaling (**G**) and DNAPK_cs_ KO and inhibition (**H**). **I-M**) M- and N-synergies discussed in the text. Column labels are names of proteins or the inhibitor used (‘i’ suffix). Proteins or inhibitors with significant CCDs (FDR_CCD_ < 0.05) are marked with an asterisk. Chromatin features are colored as in **Fig. 1C**. (**I**) ATM signaling. (**J**) Fanconi anemia complex. (**K**) SMC5/6 complex. (**L**) DNAPK_cs_ KO and inhibition. (**M**) BRCA1-A complex.

### Heterochromatin M-synergy of the FANC complex

Additionally, we found a clique that consists of Fanconi anemia (FA) proteins (FANCF, FANCM, FANCG and FANCD2) (**Fig. 3E**) and a third clique with two FA proteins together with BLM and RMI2 (**Fig. 3F**). Although FA proteins are primarily known to be involved in repair of inter-strand cross-links (*32*), they have also been implicated in MMEJ (*18*). Six out of 12 tested FA proteins show selective M-synergies with either H3K27me3 or triple heterochromatin, or both. Moreover, four additional FA proteins (FANCA, FANCB, FANCC and FANCI) showed similar trends although they individually did not pass the significance threshold (**Fig. 3J)**. These results indicate that the FA complex is an important regulator of MMEJ:NHEJ balance in heterochromatin.

### M-synergy of the SMC5/6 complex

Another complex implicated in DSB repair in heterochromatin is the SMC5/6 complex (*28, 33*). SMC5, NSE1 (NSMCE1) and NSE3 (NSMCE3) exhibit M-synergies with H3K27me3 and LMNB1. SMC6 displays similar M-synergy although it did not pass the significance threshold (**Fig. 3K**). These data indicate that the SMC5/6 complex preferentially modulates MMEJ:NHEJ balance in H3K27me3 and lamina-associated heterochromatin.

### Highlights of N-synergies

Among canonical components of the NHEJ pathway, only POLL exhibits significant N-synergy. Our data indicate that the ability of POLL to promote NHEJ is facilitated by euchromatin, particularly in transcribed regions. DNA-PKcs (PRKDC), another crucial regulator of NHEJ, showed only a weak, non-significant N-synergy pattern (**Fig. 3H**). However, treatment of cells with the DNA-PKcs inhibitor M3814 yielded a pattern that was similar but much stronger (**Fig. 3L, Fig. S6A-B**). Treatment with a potent small-molecule inhibitor is expected to have a higher penetrance than the Cas9-mediated KOs in our screen, which have incomplete efficacy (Methods and **Fig. S7**). The consistent pattern indicates that DNA-PKcs is primarily N-synergistic with transcribed parts of the genome, and to a lesser extent with triple heterochromatin. Another protein known to promote NHEJ is BD1L1 (*34*). Our data indicate that this regulatory role of BD1L1 is restricted to euchromatin (**Fig. S4E**), while it additionally shows M-synergy in triple heterochromatin (**Fig. S4F**). Also noteworthy is CAF1A (CHAF1A), the only protein that exhibits exclusive N-synergy with triple heterochromatin. CAF1A is a component of the CAF1 complex that is particularly important for nucleosome assembly in heterochromatic parts of the genome, and it has previously been found to interact with NHEJ gatekeepers KU80 and DNA-PKcs (*35*).

Other N-synergistic proteins have previously been linked to various other repair pathways, underscoring extensive cross-talk between pathways (*36*). An example is the BRCA1-A complex, which fine tunes BRCA1-mediated resection (*37, 38*). Its subunits BRCC36 (BRCC3), RAP80 (UIMC1) and BRE (BABAM2) exhibit N-synergies with euchromatic features, and ABRAXAS1 shows similar patterns but did not pass the significance threshold (**Fig. 3M**).

### Impact of CCD of BRCA2 on human cancer genomes

Furthermore, BRCA2 shows N-synergy with euchromatin (**Fig. S4A**). In this case the N-synergy may be due to local suppression of MMEJ, because suppression of MMEJ by BRCA2 has been reported (*39, 40*). To further validate these findings and to study the potential impact on genome-wide DSB repair in human cancer, we compared the genomic distribution of short deletions with either MMEJ or NHEJ signatures in BRCA2^-/-^ and BRCA2^+/+^ genome-instable tumors (data from (*41*); **Table S9-S10**; see Methods). MMEJ and NHEJ deletions are more frequent in BRCA2^-/-^ tumors compared to BRCA2^+/+^ tumors, unlike other indel signatures (**Fig. S8A**). Based on our screen results (**Fig. S4A**), we predicted that in BRCA2^-/-^ tumors − compared to BRCA2^+/+^ tumors − the log_2_MMEJ:NHEJ ratio should increase relatively more in euchromatin compared to lamina-associated heterochromatin. Indeed, this is what we observed (**Fig. 4A**). Large deletions (> 1.4kb) with MH at their break sites (which we assume to be primarily repaired by MMEJ) also showed a striking shift towards euchromatin in BRCA2^-/-^ tumors compared to BRCA2^+/+^ tumors (**Fig. S8D**), consistent with N-synergy of BRCA2 with euchromatin (**Fig. 4B**). This result illustrates that CCD of a repair protein can directly impact the type of mutations that accumulate in different chromatin contexts in tumors.

**Fig. 4.**
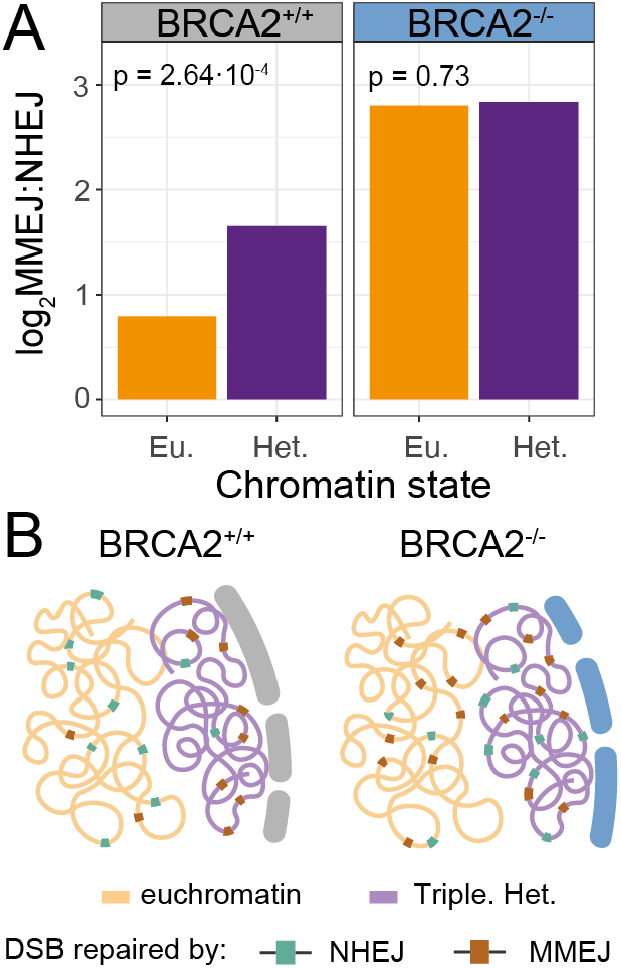
Impact on mutation distribution in cancer genomes. (**A**) Total MMEJ and NHEJ signature indel log_2_ ratio in euchromatin (Eu.) and constitutive lamina-associated heterochromatin (Het.) of BRCA2-positive (BRCA2^+/+^, n = 22) and BRCA2-deficient tumors (BRCA2^-/-^, n = 41). p-values are calculated by Fisher’s exact test applied to the total count of each deletion signature per tumor type (**Fig. S8B-C**). (**B**) Cartoon illustrating BRCA2 N-synergy.

### Overall interpretation and implications of the study

Our data uncover a much more complex network of regulatory interplay between repair proteins and chromatin components than previously thought (*42*). Generally, the effect sizes of CCDs of individual proteins that we uncovered here are modest (typically do not exceed ∼50%, see Methods, **Fig. 2F**). However, redundancies may obscure effects of individual KOs, and penetrance of the KOs in our screen is incomplete (∼65%; see Methods, **Fig. S7**), causing underestimation of the effect sizes. Considering the large number of proteins exhibiting CCDs, their collective effect is likely to be substantial. Most likely, other DNA repair pathways that we did not probe are also subject to extensive CCDs. Multiplexed IPR screens as described here help to uncover these regulatory networks and provide a foundation for further exploration of the underlying molecular mechanisms.

## ACKNOWLEDGEMENTS

We thank the NKI Genomics and Research High Performance Computing core facilities for technical support; members of our laboratories and Titia Sixma for inspiring discussions and helpful comments; Jacqueline Jacobs, Heinz Jacobs and Jeroen van den Berg for help with KO library gene set curation.

## Author Contributions

X.V., A.G.M, R.S, B.v.S, R.H.M conceived and designed the study. A.G.M, B.M and X.V. performed experiments. X.V., C.L. and M.S. performed data analysis. B.v.S, R.H.M and R.L.B supervised the study. X.V., B.v.S and R.H.M. wrote the manuscript with input of all authors.

## Data and materials availability

All data shown in this paper have been deposited in the Short Read Archive (BioProject no. PRJNA882344). All code to analyze the data and create the figures is available on GitHub (https://github.com/vansteensellab/CCD_repair_protein_project).

## SUPPLEMENTARY MATERIALS

Materials and Methods

Figs. S1 to S8 Scheme 1 to 6

Tables S1 to S10

References (43-57)

## MATERIALS AND METHODS

### A. EXPERIMENTAL PROCEDURES Cell line and culture conditions

We used the clonal cell line K562#17 DSB-TRIP clone 5 (*6*), which is a genetically modified monoclonal human K562 cell line (ATCC). This cell line stably expresses Shield1-inducible DD-Cas9 and additionally carries 19 uniquely barcoded integrated pathway reporters (IPRs) in precisely mapped genomic locations (**Table S6**). Cells were cultured in RPMI 1640 (GIBCO) supplemented with 10% fetal bovine serum (FBS, Capricorn Scientific) and 1% penicillin/streptomycin. Cells were regularly checked to be free of mycoplasma.

#### Design of KO gRNA library

We designed an arrayed CRISPR/Cas9 KO gRNA library (KO gRNA library, in short) which targeted a total of 519 genes encoding proteins previously linked to DNA repair. The list of proteins was based on the Gene Ontology term GO:0006302 (double strand break repair), supplemented with a manually curated list. The crRNA library was generated by Integrated DNA Technologies (IDT) and contained 4 gRNAs per gene (**Table S1**). The individual crRNAs were delivered in a lyophilized RNA form and were diluted in Duplex Buffer (DB, IDT cat. no. 11-01-03-01) to a stock concentration of 100 μM. Finally, we pooled crRNA targeting the same gene to a single well in a final concentration of 5 μM per crRNA.

#### Screen procedure

##### Overview

We performed the screen in 96-well format. It consisted of the following key steps:

- Day 1: Induction of Cas9 expression and transfection with gRNAs to disrupt 519 individual genes (**scheme 1**)
- Day 5: Passaging of cells; quality checks of liquid handling and transfection efficiency (**scheme 2**)
- Day 6: Second transfection: induction of DSBs in the IPRs (**scheme 3**)
- Day 9: Lysis of cells (**scheme 4**)
- Downstream processing: PCR amplification and sequencing of the barcoded IPRs (**scheme 5**)

##### Liquid handling

Steps in the procedure were performed in a semi-automated fashion either with MicroLab STAR liquid handler (Hamilton Company, **blue in scheme 1-5**), Multidrop™ Combi Reagent Dispenser (ThermoFisher, **green in scheme 1-5**) or manually (**grey in scheme 1-5**).

##### Day 1: Induction of Cas9 and transfection of KO gRNA library (**scheme 1**)

Eight hours before the KO gRNA library transfection, we diluted clone 5 to a final concentration of 85,000 cells/ml with medium containing 500 nM Shield-1 (Aobious cat. no. AOB1848) to stabilize DD-Cas9 protein. As first step in the KO gRNA library transfection, we diluted 20 μM tracrRNA (IDT cat. no. 1072534) stock concentration to 800 nM in Duplex Buffer (DB, IDT cat. no. 11-01-03-01) in a final volume of 24 μl. Next, we pipetted 1μl crRNA of KO gRNA library (stock at 20 μM in DB) or controls to its appropriate position in the gRNA plate (orange, scheme 1). Plates 1 to 5 in the screen included 88 KO gRNAs and 8 controls wells: four mock KO controls (crRNA was omitted), one POLQ KO gRNA control (used as a positive control, sequences in **Table S2)**, one editing control (transfected with LBR2 gRNA (*43*)) and one pipetting control. In the pipetting control, 1 μl phenylarsine oxide (PAO Sigma-Aldrich, cat. no. P3075, stock concentration of 10 mM) was pipetted instead of 1 μl crRNA. 10 μM of PAO is enough to kill K562 cells, so we used visual inspection of cell death at day 5 to check if the KO gRNA library pipetting step was successful. Plate 6 included nine additional mock KO controls, making a total of 33 per replicate. In parallel, we diluted DharmaFect #4 (Horizon Discovery, cat. no. T-2004-03) lipofectamine to 2.66% (0.4 μl in 15 μl) with Optimem (Gibco, cat. no. 31985070). After 5 minutes incubation at room temperature, 15 μl of diluted DharmaFect #4 was pipetted in the three empty 96-well V-bottom plates (Thermo Fisher, cat. no. 11816003) and 5 μl of the 800 nM coupled gRNA. This mix was incubated for 15 minutes at room temperature and subsequently 15,000 clone 5 cells were dispensed per well (180 μl of clone 5 cells in 500 nM Shield-1). This procedure was repeated six times, once for each different KO gRNA library plate. We note that every new batch of DharmaFect #4 lipofectamine was tested and the cell:lipofectamine ratio was adapted for optimal transfection efficiencies. The reagent quantities described above are representative of the concentrations used in the screen.

**Scheme 1:**
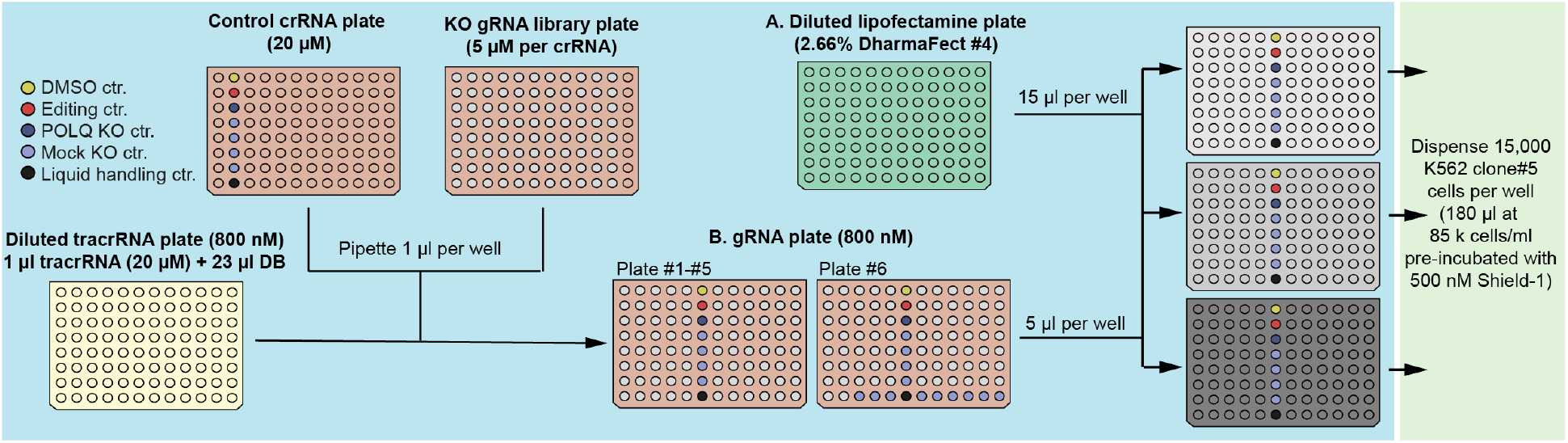
Day 1 procedure − KO gRNA library transfection (repeated 6 times). 24 μl of 800 nM tracrRNA (yellow) was pipetted to the final crRNA:tracrRNA plate (orange). Then, 1 μl of KO gRNA library or control plate added to screening plate position. Next, Diluted DharmaFect #4 was pipetted to lipofectamine plate (green). 15 μl of the lipofectamine was pipetted into three 96-well plates (three replicates) and 5 μl of the crRNA:tracrRNA mix added later. With the cell dispenser, 15,000 clone5 cells per well were dispensed (green).

#### Day 5: passaging of cells and quality checks (**scheme 2**)

Four days after KO gRNA library transfection, we split transfected cells 1:10 with fresh medium supplemented with 500 nM Shield-1. At this step, we harvested editing control wells for TIDE analysis and we visually inspected cell death in the pipetting control wells. We used these two quality controls to assess if specific plates should be discarded or kept for the following steps. We repeated this process for every plate in the screening.

TIDE (*44*) was used to monitor the editing efficiency prior to high-throughput sequencing, as follows. Editing control wells were harvested and cells were lysed with 30 μl DirectPCR lysis buffer (Viagen cat. no. 301-C) supplemented with 1 mg/ml proteinase K (Bioline, cat. no. BIO-37084) by incubating them at 55 °C for at least 2 hours up to overnight, followed by heat inactivation for 45 min at 85 °C. To monitor the CRISPR editing frequency, we used primers spanning the endogenous LBR2 target site as previously reported (*43*). PCR was performed using 10 μl MyTaq Red mix (Bioline, cat. no. BIO-25044), 1 μM of each TAC0017 and TAC0018 primers, 2 μl of cell lysate and up to 20 μl of water. PCR conditions for TIDE analysis are the following ones: 1 min at 95 °C followed by 28 cycles of 15 s at 95 °C, 15 s at 58 °C and 30 s at 72 °C and a final extension of 1min at 72 °C. The excess of PCR primers was degraded by EXOSAP treatment as follows. For each 10 μl of PCR reaction, 0.125 μl of Shrimp Alkaline Phosphatase (1 U/ml; New England Biolabs, cat. no. M0371S), 0.0125 μl Exonuclease I (20 U/ml; New England Biolabs, cat. no. M0293S) and 2.36 μl of water were added. Samples were incubated at 37 °C for 30 min and heat inactivated for 10 min at 95 °C. Next, 5 μl of EXOSAP-treated PCR mix was Sanger sequenced with 5 μl of TAC0017 primer at 5 μM concentration by Macrogen (EZ-seq). The resulting Sanger sequence traces were analyzed using the TIDE algorithm (*44*) to determine the editing efficiency.

**Scheme 2:**
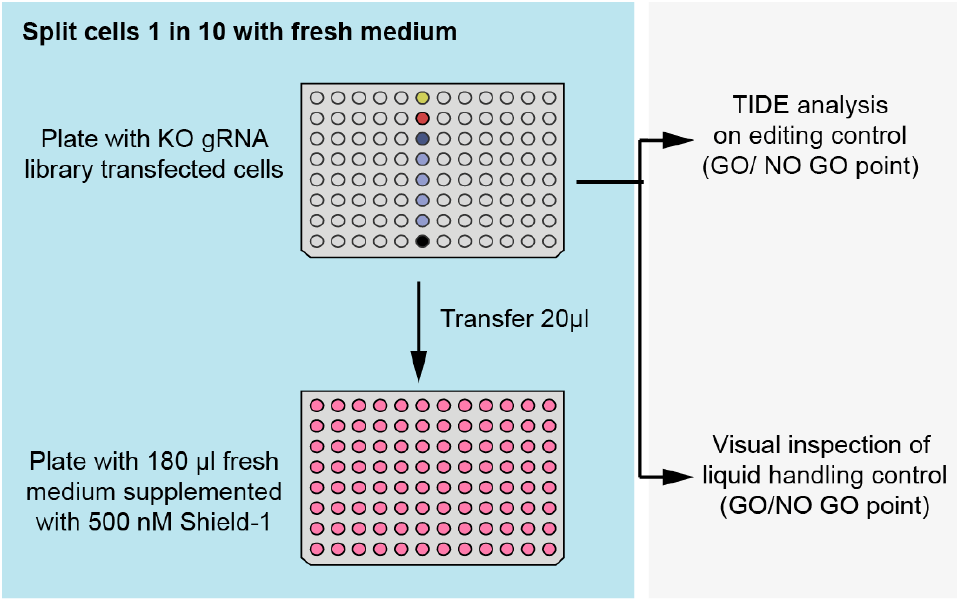
Day 5 procedure - Passaging of cells and harvesting of control samples (repeated 18 times). Screening plates were split 1:10 with fresh medium. Editing control was harvested for TIDE analysis and the pipetting control was checked visually.

##### Day 6: induction of DSBs in IPRs by transfection with LBR2 gRNA (**scheme 3**)

We manually mixed LBR2 crRNA (crRNA targeting DSB-TRIP reporters) with tracrRNA at a final concentration of 800 nM in DB and diluted DharmaFect in Optimem (2.66% concentration). Then, we pipetted 15 μl of diluted lipofectamine into six empty 96-well plates and added 5 μl of LBR2 crRNA:tracrRNA. After incubating this mix for 15 min at room temperature, we added 180 μl KO cells in arrayed format. We repeated this procedure for each replicate independently with freshly prepared LBR2 gRNA and DharmaFect #4 mixes.

**Scheme 3:**
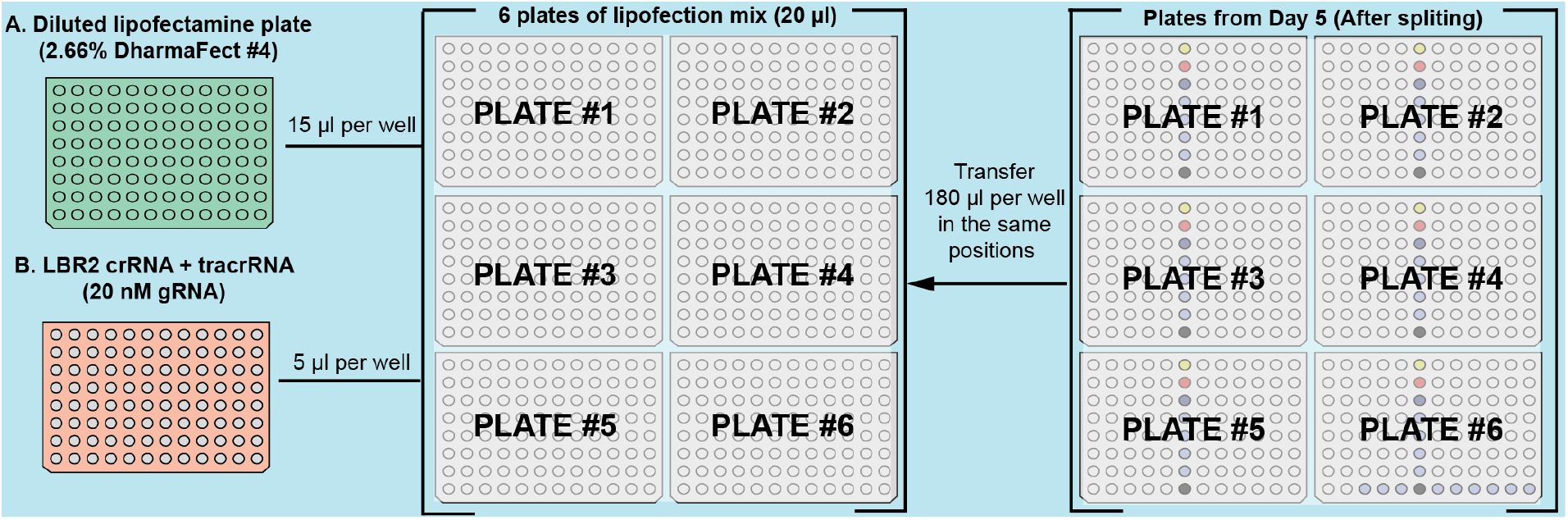
Day 6 procedure - LBR2 gRNA transfection (repeated 3 times). DharmaFect #4 and LBR2 crRNA:tracrRNA were manually diluted. Then, 15 μl DharmaFect #4 and 5 μl LBR2 crRNA:tracrRNA (B on A) was combined into each well of 6 empty 96-well plates. 15 minutes after mixing, 180 μl of cells, split on day 5, were transferred into the transfection mix plate.

##### Day 9: cell lysis and quality controls (**scheme 4**)

Three days after LBR2 gRNA transfection we harvested the screening plates. To do so, we centrifuged 96-well plates to pellet the cells (1500 rpm for 5 min). Then, we removed the supernatant and pipetted 30 μl of DirectPCR lysis buffer supplemented with 1 mg/ml proteinase K on the cell pellets. After a couple of pipetting cycles to mix cells with lysis buffer, we transferred the cell lysate to an empty 96-well PCR plate (ThermoFisher, cat. no. AB0900). Cells were lysed overnight at 55°C in a thermocycler, and proteinase K was subsequently inactivated for 45 minutes at 85°C.

To monitor CRISPR editing efficiency of the second transfection, we performed TIDE analysis with cell lysate from a random well from each plate, as described above (Day 5).

**Scheme 4:**
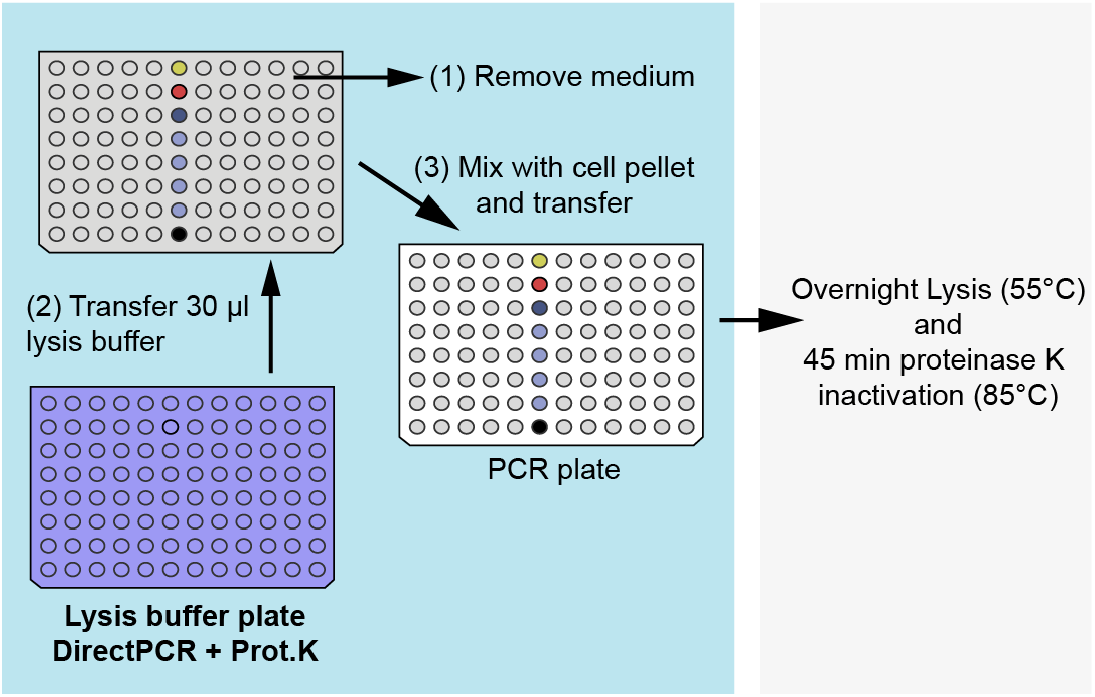
Day 9 procedure - Screening plate harvesting (repeated 18 times). Harvesting was performed in three steps. After cells were pelleted by centrifugation, 170 μl supernatant was removed (grey, 1); lysis buffer was added (purple) and mixed with cell pellets (grey, 2); and the cell lysate was transferred to empty PCR plates (white, 3). Cells were lysed overnight at 55°C and proteinase K was subsequently heat inactivated for 45 min at 85 °C.

##### Screening replicates

The screening was performed twice, more than a month apart. Each time the screening was performed in three replicates with independent transfection mixes. Three out of six replicates were discarded because of technical reasons such as wrong liquid handling, unsuccessful transfection (at least ∼50% editing in the editing efficiency control) or problems during sample processing. One replicate from the first screen and two replicates from the second screen passed the quality controls. We refer to these replicates as replicate 1 (R1), replicate 2 (R2) and replicate 3 (R3).

##### Downstream processing: sample preparation for IPR sequencing (**scheme 5**)

For the sequencing of the IPRs (to identify indels and their linked IPR barcodes) in all screen samples, we employed a two-step PCR indexing and pooling strategy as previously described (*6*), with some adaptations. We performed the first PCR reaction (indelPCR1) with TAC0007 (indexed) and TAC0012 (non-indexed) primers with a unique TAC0007 indexed primer for each 96-well plate, and the second PCR reaction (indelPCR2) with TAC009 (non-indexed) and TAC0159 (indexed) primers with 96 different TAC0159 primers (one for each well in a 96-well plate). Pipetting was performed using the MicroLab STAR liquid handler (Hamilton Company).

IndelPCR1 and indelPCR2 were performed under similar PCR conditions with the only difference being the number of cycles. In both reactions, a denaturing step was performed for 1 min at 95 °C, low annealing temperature amplification cycles (cold cycles) for 15 s at 95 °C, 15 s at 55 °C and 15 s at 72 °C, high annealing temperature amplification cycles (hot cycles) for 15 s at 95 °C, 15 s at 70 °C and 15 s at 72 °C and a final extension of 2 min at 72 °C.

**Scheme 5:**
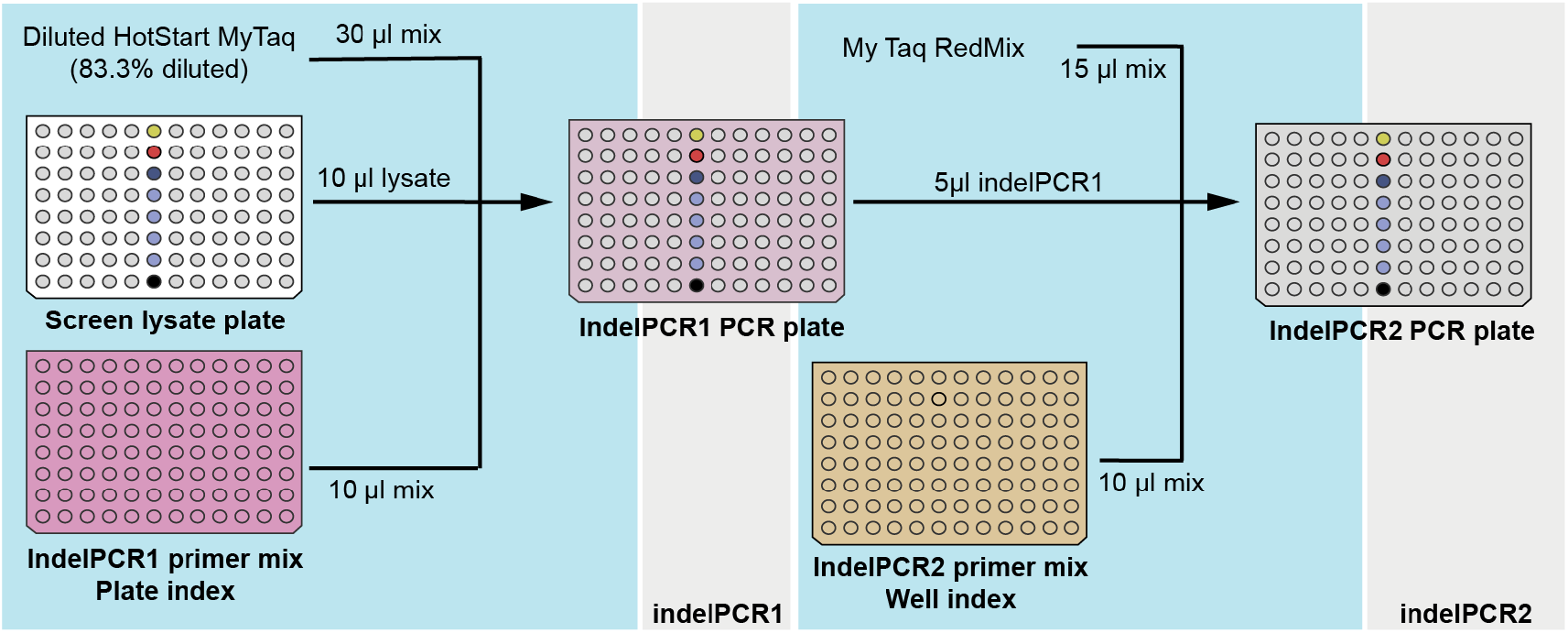
Screening sample preparation for sequencing. PCR amplification of indel and barcode of each IPR in all screening samples was performed in two steps: indelPCR1 and indelPCR2. Pipetting was performed with the liquid handler (blue) and PCRs with a ThermoCycler (grey). IndelPCR1 was pipetted from three source plates: Diluted HotStart MyTaq Red Mix (5 parts of mix with 1 part of H2O), screening cell lysates (white) and indelPCR1 primer mix (purple). A different indexed primer was used per plate. IndelPCR2 was also pipetted from three source plates: MyTaq Red mix, indelPCR1 PCR plate (light purple) and indelPCR2 primer mix (gold). A different indexed primer was used for each well.

We performed indelPCR1 with 10 μl of cell lysate from the screening plates, 30 μl of 86.6% (5:1) diluted MyTaq HotStart Red Mix in water (Bioline, cat. no. BIO-25048) and 10 μl of 1 μM of each primer (TAC0007 and TAC0012 final concentration of 200 nM) for 4 cold cycles and 9 hot cycles. Then, we performed indelPCR2 with 5 μl of indelPCR1 product, 15 μl of MyTaq Red mix (Bioline, cat. no. BIO-25044) and 10 μl of 500 nM of each primer (TAC0009 and TAC0159, final concentration of 166 nM) for 3 cold cycles and 8 hot cycles.

Next, we pooled indelPCR2 products per plate in equal volumes and DNA was purified with cleanPCR (CleanNA cat. no. CPCR-0050) beads at a 0.8:1 beads:sample ratio. Ten μl of each pool was run on a 2% agarose gel for visual inspection, and DNA concentration was quantified by Qubit DNA dsHS Assay Kit (Invitrogen, cat. no. Q32851). Equimolar concentrations of DNA per plate were pooled and the resulting product run on a 2% agarose gel. The PCR amplicon band was cut from the gel and isolated by PCR Isolate II PCR and Gel Kit (Bioline, cat. no. BIO-52060) and lastly bead-purified. Resulting preparations were sequenced on a NextSeq MID with single-ended 150 bp reads with ∼25 % of PhiX spike-in.

#### Chromatin context effects assessed with inhibitors

For DNAPK and ATM we also determined CCDs by using specific small-molecule inhibitors. The experimental design was similar to the KO screen setup, with the following modifications:

##### Plasmid transfection

To induce DSBs, we introduced LBR2 gRNA into the cells by plasmid nucleofection instead of RNA transfection. For this purpose, we resuspended one million K562 clone 5 cells in 100 μl transfection buffer (100 mM KH2PO4, 15 mM NaHCO3, 12 mM MgCl2, 8 mM ATP, 2 mM glucose (pH 7.4)) (*45*). Then, we added 12 μg of either gRNA-containing LBR2 plasmid or GFP-expressing control plasmid. Cells were electroporated in an Amaxa 2D Nucleofector (T-016 program). 24 hours post-nucleofection, we assessed transfection efficiency by visual observation of GFP-positive cells. This GFP sample was later used as non-targeted control.

##### Inhibitor treatment

Eight hours after nucleofection, we added 500 nM Shield-1 (Aobious) to stabilize DD-Cas9 protein. Together with Shield-1, we added inhibitors of either DNAPK (M3814, final concentration 1 μM from a 1 mM stock in DMSO, MCE cat. no. HY-101570), ATM (KU5593, final concentration 10 μM from a 10 mM stock in DMSO, Calbiochem cat. no. #118500), and DMSO-only vehicle controls (1:1000, Sigma cat no. D4540).

##### Indel library preparation

72 hours after DD-Cas9 stabilization, we harvested the cells, performed genomic DNA (gDNA) extraction with the ISOLATE II genomic DNA kit (Bioline, BIO-52067) and diluted DNA to 50 ng/μl. Indel sequencing libraries were prepared as described for the screen but with minor changes as follows. We performed indelPCR1 with 200 ng of gDNA as input (4 μl of 50 ng/μl concentrated sample) and 200 nM of each primer for 4 cold cycles and 8 hot cycles. Then, we performed indelPCR2 with 5 μl indelPCR1 product and 166.6 nM of each primer for 1 cold cycle and 13 hot cycles. We pipetted both PCR reactions manually. We pooled samples in equimolar ratios and prepared them for sequencing as described for the screen. Samples were sequenced in a MiSeq Nano (Illumina) with 10 % of PhiX spike-in. We performed this experiment in three independent biological replicates.

##### Data analysis

We analyzed the CCDs of inhibitors as performed for the screen data (see data processing section below), with slight modifications. We tested the significance of the perturbation by means of a Student’s t-test instead of a z-test. We used this test because here each replicate includes only a single control sample. Everything else was performed as described for the screen data.

### B. DATA PROCESSING AND COMPUTATIONAL ANALYSES

#### Processing and statistical analysis of screen data

**Scheme 6:**
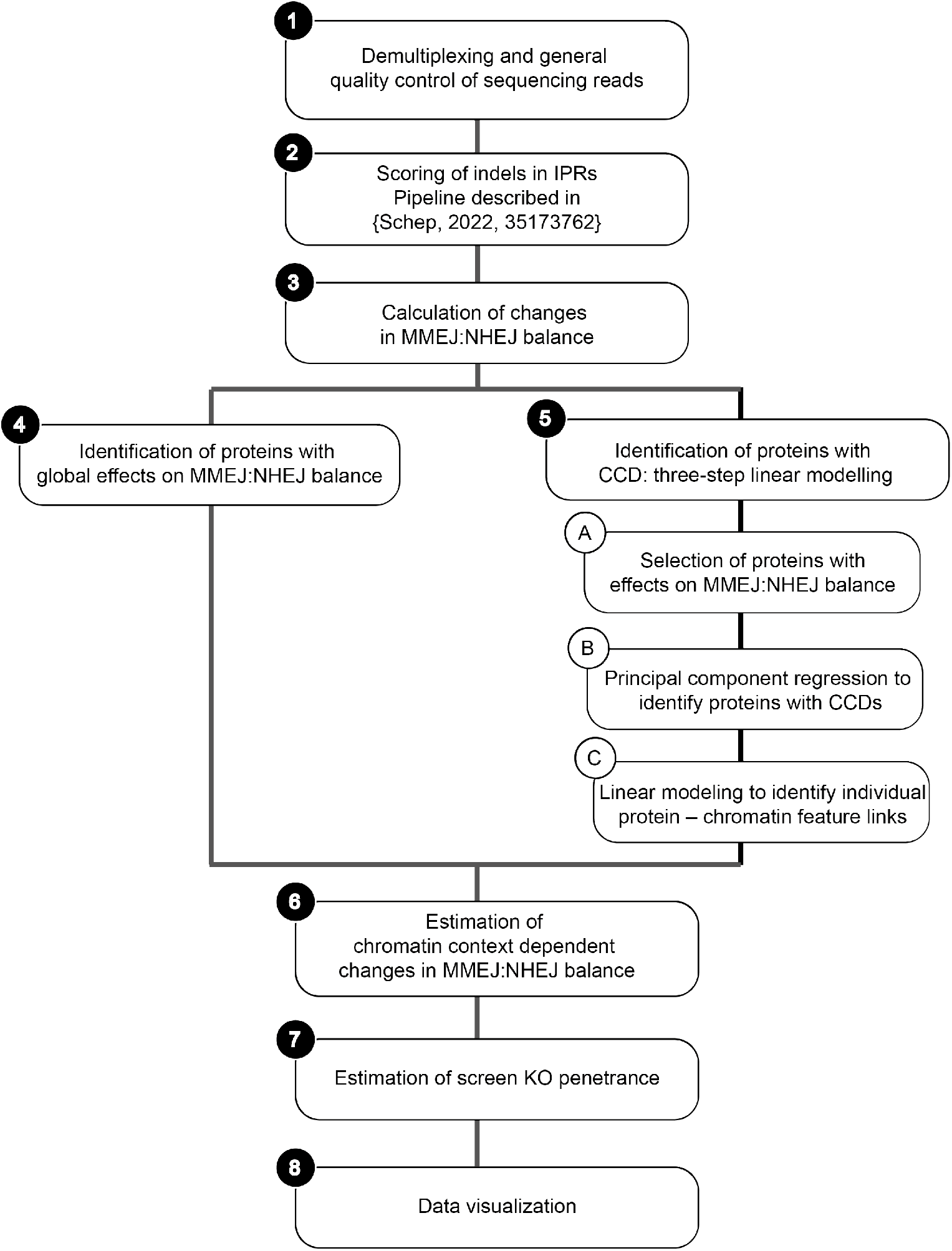
Key steps in the data processing and computational analysis workflow.

##### 1. Demultiplexing and general quality control of sequencing reads

Demultiplexing of the sequencing reads was done based on indices added in the IndelPCR1 (plate index) and IndelPCR2 steps (well index) and each file contains the reads from a single well in the screen. We refer to these as sample throughout this section. Demultiplexed sequencing data is available in the Sequence Read Archive (https://www.ncbi.nlm.nih.gov/sra; BioProject no. PRJNA882344). An overview of obtained read numbers is provided in Table S3.

##### 2. Scoring of indels in IPRs

Scoring of indels and linking to their IPR barcodes in the sequence reads was done using a previously reported computational pipeline (*46*). In short, for each sequence the barcode was extracted and the indel state was classified. As documented previously (*6, 43*), a single-nucleotide insertion was assumed to be created by NHEJ repair (NHEJ_ins_), a seven-nucleotide deletion was assumed to be created by MMEJ repair (MMEJ_del_), and the absence of indel was assumed to be intact DNA (uncut or perfectly repaired). In the downstream analysis, we use the number of NHEJ_ins_ reads, MMEJ_del_ reads and intact reads. The median and 95% CI number of reads per replicate are summarised in Table 5.

**Table S3:**
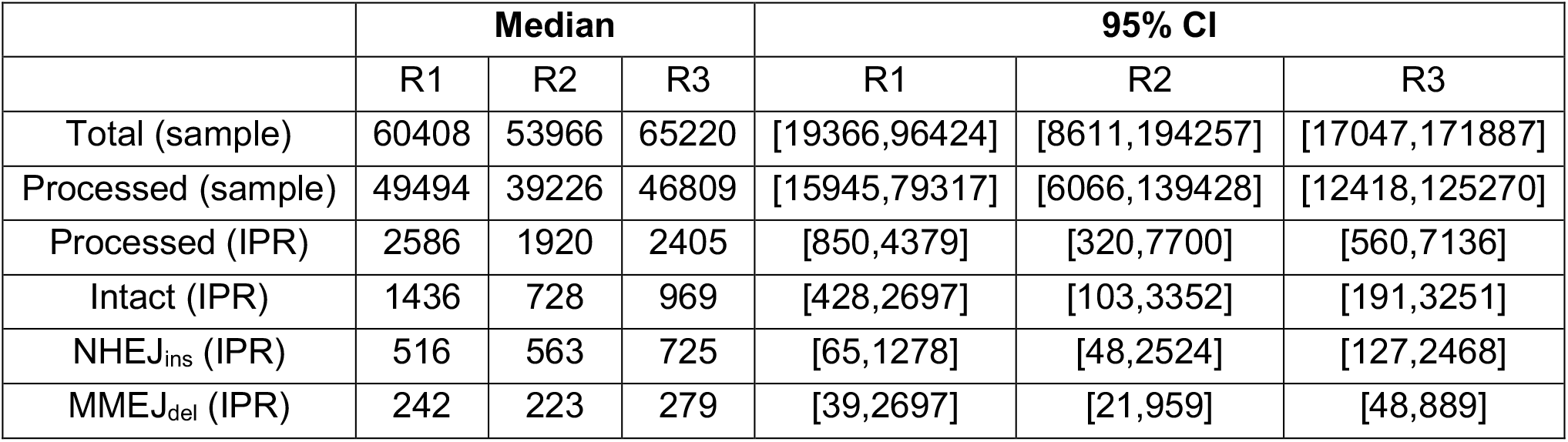
Overview of sequencing read numbers per sample in the screen

##### 3. Calculation of changes in MMEJ:NHEJ balance

After indel scoring, we calculated the editing efficiency (Formula 1) and log_2_ MMEJ:NHEJ balance (Formula 2) for each individual IPR in every sample. Next, we filtered out data based on two parameters: low read numbers and low editing frequency. First, we discarded IPRs which had less than 30 reads with either NHEJ_ins_ or MMEJ_del_ per sample. Second, we discarded samples with an average editing efficiency lower than 25% per sample. After these filtering steps, 531 samples from R1, 541 samples from R2 and 555 samples from R3 were retained, with an average of 2.92 replicates and 18.98 IPRs per well.

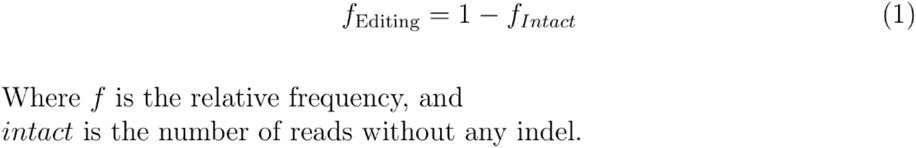

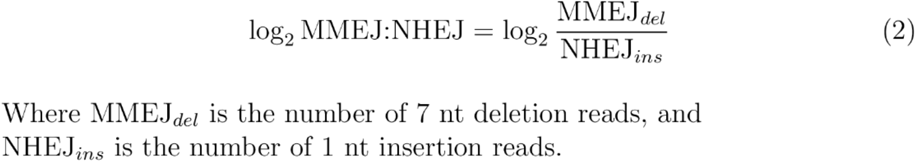

Then, we divided samples into three categories depending on the gRNA they received in the first transfection (Day 1): mock KO controls, POLQ KO controls and KO gRNA library samples. We checked the reproducibility of the log_2_MMEJ:NHEJ balance between replicates (**Fig. S1A-C**). Next, we computed for each IPR the log_2_ fold change in MMEJ:NHEJ balance (Δlog_2_MMEJ:NHEJ) as a consequence of each KO (Formula 3) and averaged three replicates.

A negative Δlog_2_MMEJ:NHEJ score implies either reduced MMEJ or increased NHEJ activity (at the tested IPR) due to the KO of the tested protein. Our assay cannot discriminate between these two possibilities, as we cannot measure the individual pathway activities − only the balance (*6, 43*). Likewise, a positive Δlog_2_MMEJ:NHEJ score implies either increased MMEJ or decreased NHEJ activity (at the tested IPR). For simplicity, we refer to a negative Δlog_2_MMEJ:NHEJ score as: the tested protein (when present) *favors MMEJ*; and we refer to a positive Δlog_2_MMEJ:NHEJ score as: the tested protein (when present) *favors NHEJ*.

The POLQ KO samples provide an indication of the dynamic range of Δlog_2_MMEJ:NHEJ scores that may be expected (**Fig. S1D**), because POLQ is essential for MMEJ (*47, 48*). On average, POLQ KO samples showed a Δlog_2_MMEJ:NHEJ score of -1.58 log_2_ units across all IPRs, i.e., a ∼3.0-fold reduction in MMEJ:NHEJ balance. As most proteins are not absolutely essential for either MMEJ or NHEJ, the dynamic range of the Δlog_2_MMEJ:NHEJ scores may be expected to be less than the score observed for POLQ. Indeed, this is the case (**Fig. S1D**).

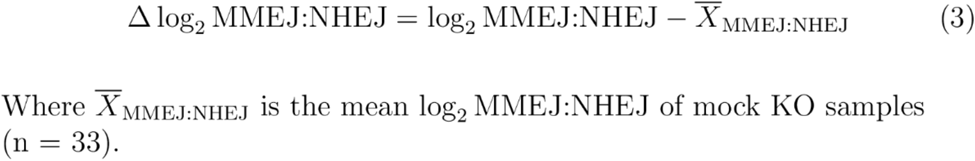

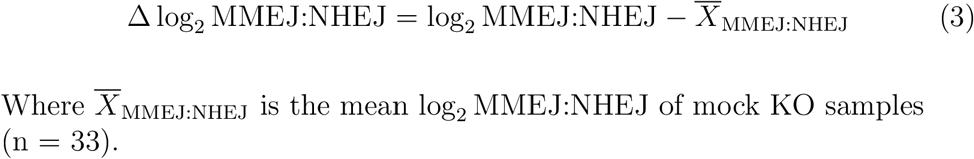

We used the Δlog_2_MMEJ:NHEJ scores throughout this work as a metric of the contribution of each protein to the MMEJ:NHEJ balance. For global MMEJ:NHEJ contribution of proteins, we computed the average Δlog_2_MMEJ:NHEJ over all 19 pathway reporters (**Section 4 in data analysis workflow**). When calculating chromatin context dependencies (CCDs), the Δlog_2_MMEJ:NHEJ of each IPR-KO combination was used (**Section 5 in data analysis workflow**).

##### 4. Identification of proteins with global effects on MMEJ:NHEJ balance

To assess the global effect of proteins on MMEJ:NHEJ balance (**Fig. 1B & Fig. 2**), we computed for each KO the mean Δlog_2_MMEJ:NHEJ over all 19 IPRs. Next, to identify proteins that significantly *favor MMEJ* or *favor NHEJ* independently of the chromatin state, we tested whether the mean Δlog_2_MMEJ:NHEJ was different than zero by a Student’s t-test followed by Benjamini-Hochberg multiple-testing correction of p-values. We called proteins to globally *favor MMEJ* (mean Δlog_2_MMEJ:NHEJ < 0) or globally *favor NHEJ* (mean Δlog_2_MMEJ:NHEJ > 0) with an estimated false-discovery rate (FDR) < 0.001.

##### 5. Identification of proteins with CCD: three-step linear modelling

###### a. Initial selection of proteins with any effect on MMEJ:NHEJ balance

To filter for proteins with any effect on MMEJ:NHEJ balance, we calculated the z-score log_2_MMEJ:NHEJ for each 19 IPR in 519 KO gRNA samples (total of 9861) using the 33 mock KO gRNA samples (total of 627) to empirically estimate null-distributions. First, we fitted a normal distribution through the mock KO log_2_MMEJ:NHEJ scores (n = 33) for each IPR and replicate separately (example IPR in **Fig. S2A**). Next, we standardized log_2_MMEJ:NHEJ scores of each sample using the mean and standard deviation of the fitted distributions (Formula 4) (example IPR in **Fig. S2B**). Finally, we combined the z-scores of the three independent replicates by Stouffer’s method (Formula 5) (**Fig. S2C**). After this transformation, 24.5% KO - IPR combinations (n = 2420) had an absolute z-score >1.96, compared to only a 4.3% of mock KO - IPR combinations (n = 27).

We retained a KO if the absolute z-score was >1.96 in at least 2 out of 19 IPRs. A total of 352 KOs passed this filter. Of the 33 mock KO samples four passed the same criteria, suggesting an empirical FDR of 12%. Note that further filters are applied below for additional stringency. From the 352 proteins that passed this filter, 296 *favor MMEJ*, 47 *favor NHEJ* and 9 had mixed effects, i.e they *favor MMEJ* in some IPRs and *favor NHEJ* in others.

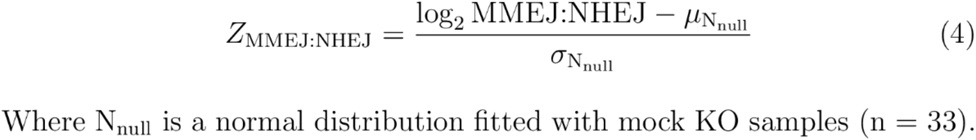

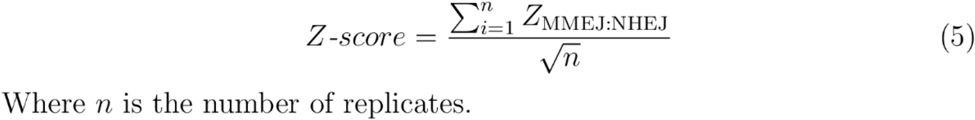

Where *n* is the number of replicate.

###### b. Principal component regression

Next, among the remaining 352 proteins, we identified proteins with significant chromatin context dependencies (CCDs) across the entire set of 25 chromatin features. Because of the strong covariation among most chromatin features, we did this by principal component (PC) regression. This consists of dimension reduction using standard principal component analysis, followed by linear regression on the main PCs (**Figure S3A**). This approach provides substantial robustness and avoids identification of fortuitous correlations with single chromatin features.

The sources of all chromatin feature tracks are summarized in **Table S2**. Each of these 25 tracks was z-normalized. Z-scores were calculated as the log_2_ fold-difference of the signal over control (matching controls as provided by the respective studies) in 2kb bins centered around each IPR insertion site. These values were subsequently converted into z-scores using the mean and standard deviation of the chromatin feature signal in the TRIP pools, as previously done (*6*).

We then first assessed the number of PCs needed to explain most of the variance in the chromatin data of the 19 IPRs. For this we used *pls* package (version 2.8-1) in *R*. We selected the first three PCs, which together account for 76% of the variance. Adding a fourth PC to the model would only increase the explained variance by 6% (**Fig. S3B**). Closer inspection of the first three PCs revealed that each PC explained biologically relevant differences in chromatin contexts: PC1 mainly explained the difference between euchromatin and heterochromatin, PC2 mainly explained differences between heterochromatin types (Triple heterochromatin vs. H3K27me3) and PC3 mainly explained differences between euchromatin types (Enhancer/promoters vs. transcription) together with replication timing (**Fig. S3C**). Then, for each of the 354 KO and 4 mock samples, we constructed a linear model based on three PCs to predict the Δlog_2_MMEJ:NHEJ scores. To assess the accuracy of this fit, we computed the p-value of the correlation between predicted Δlog_2_MMEJ:NHEJ scores and measured Δlog_2_MMEJ:NHEJ scores for each of the samples. After Benjamini-Hochberg correction of these p-values for multiple testing, 89 protein KOs and 1 mock KO passed the significance threshold at FDR cutoff 0.05.

###### c. Linear modeling to identify individual protein − chromatin feature links

Finally, to identify the individual chromatin features that contribute to the CCDs, we fitted linear correlations between Δlog_2_MMEJ:NHEJ of 89 proteins with significant CCDs and each of the 25 individual chromatin features (total of 2225). Based on this calculation, we identified individual protein − chromatin feature pairs with N-synergies, M-synergies or no synergies as follows:

A protein − chromatin feature pair is defined to have <u>N-synergy</u> when the protein *favors NHEJ* according to section *5a* in the data analysis workflow, and the linear fit has a positive slope (**Fig. S4A**). This positive slope implies that the ability of the protein to shift the balance towards NHEJ increases with increasing levels of the chromatin feature. It is also possible that a protein that *favors NHEJ* according to step *5a* shows a negative slope (**Fig. S4B**). However, such a negative correlation is likely to reflect an indirect effect. For example, IPRs with high H3K4me3 signals often exhibit low H3K9me3 signals and vice versa. Because the vast majority of molecular interactions in chromatin have so far been explained by the presence of a chromatin feature (e.g., a certain histone modification) rather than the absence of a chromatin feature, we focus on positive slopes for N-synergy and reject negative slopes as likely reflecting indirect correlations.

Conversely, we define a protein − chromatin feature pair as having <u>M-synergy</u> when the protein *favors MMEJ* (according to step *5a*) and the linear fit has a negative slope (**Fig. S4C**). Here, the negative slope implies that the ability of the protein to shift the balance towards MMEJ increases with increasing levels of the chromatin feature. Again, weaker effects with increasing levels of the chromatin feature (in this case a positive slope; **Fig. S4D**) are most likely due to indirect correlations, and thus not considered to be M-synergy.

By these criteria, a few proteins showed both M- and N-synergy, with different chromatin features (**Fig. S4E-F**). We used the slope of the linear fits of M- or N-synergistic pairs as a measure of the synergy (synergy score). This score is set to 0 for protein − feature pairs without synergistic interactions as defined above. Of the 89 proteins with significant CCDs, 73 have M-synergies, 14 have N-synergies and 2 have mixed synergies.

Additionally, we fitted similar linear models for protein − chromatin feature combinations for the remaining 263 proteins that modulate the MMEJ:NHEJ balance (step *5a*) but did not pass the CCD significance threshold (step *5b*). We highlight some of these proteins in the main text, but always in connection with proteins with significant CCDs (**Fig. 4B-D**).

##### 6. Estimation of chromatin context dependent MMEJ:NHEJ balance changes

As stated above, the synergy score is the slope of the linear fit between Δlog_2_MMEJ:NHEJ and a chromatin feature. The predicted effect size of a chromatin feature on the MMEJ:NHEJ balance (i.e., the dynamic range of Δlog_2_MMEJ:NHEJ values across the entire genome, from the lowest to the highest level of the chromatin feature) not only depends on this slope, but also on the dynamic range of levels of this chromatin feature. To estimate this effect size, we first approximated this genome-wide dynamic range of each chromatin feature from the chromatin scores of 2,150 previously characterized randomly integrated IPRs (*6*) (**grey distributions in Fig. S5A**) as the difference between the bottom 0.5% and top 0.5% (**Fig. S5A**). We then multiplied this difference with the synergy score, resulting in a rough estimate of the genome-wide CCD Δlog_2_MMEJ:NHEJ. For global and CCD Δlog_2_MMEJ:NHEJ comparisons of each proteins (**Fig. 2F**), we selected the maximum estimated CCD Δlog_2_MMEJ:NHEJ of each protein. For this figure we classified proteins with CCDs into proteins with only CCDs effects (FDR_CCD_ < 0.05 & FDR_global_ ≥ 0.001) and proteins with both CCDs and global effects (FDR_CCD_ < 0.05 & FDR_global_ < 0.001).

##### 7. Estimation of screen KO penetrance

The effect sizes calculated above are likely to be underestimates, because the KO efficiencies after transfection of the gRNAs (Day 1) are expected to be less than 100%. Because we could not measure these efficiencies for all KOs directly (which would require gene-specific PCR for each KO), we obtained an approximate estimate as follows. We assumed that Day 1 transfections were equally efficient as the Day 6 transfections. From the latter, we calculated the mean editing efficiency (Formula 1) of IPRs in transcriptionally active chromatin (n = 8). We focused on IPRs in transcriptionally active chromatin because they are more representative of the chromatin type that most gRNAs in the KO library target. We considered that a reporter is embedded in transcriptionally active chromatin when at least one of the transcription-related features TTseq, H3K36me3, POL2AS2 or POL2 had a chromatin z-score higher than 0.5 (8 pathway reporters marked by black bar in **Fig. S7A**). Then, we calculated the average editing frequency of the mock transfected samples (n = 33) for each IPR and replicate (**Fig. S7B**). The results suggest that the editing efficiency was in the range of 40-80%. This estimate is consistent with the efficiency of editing of the LBR gene with gRNA LBR2 after the first transfection, as measured on Day 5 (see above), which was in the range 47.5%-70% for the three screen replicates. However, we note that this estimation does not take into account the percentage of in-frame indels created by CRISPR/Cas9 or other gene editing products that do not lead to a protein KO.

##### 8. Data visualization

We visualized CCDs of proteins as a heatmap (**Fig. 2D**) and as a Uniform Manifold Approximation and Projection (UMAP) plot (**Fig. 3C**). In the heatmap, we hierarchically clustered the synergy scores of every protein − chromatin feature pair using the “ward.D” algorithm in the *pheatmap* package in R (version 1.0.12). The hierarchically clustered dendrogram (**Fig. 2D**) was divided into four groups to highlight the main clusters observed in the heatmap. The UMAP was calculated with the *umap* package (version 0.2.8.0) and two UMAP dimensions plotted as a scatterplot.

#### Comparison to protein-protein interaction data

To assess if physically interacting proteins tend to have similar CCDs, we computed cosine similarities of synergy scores (Formula 6) between physically interacting protein pairs and compared them to the synergy scores expected by random chance. First, we computed the cosine similarity matrix for all proteins with significant CCDs with the *lsa* package (version 0.73.3). For this we compared the 25 synergy scores for each protein. We decided to use the cosine distance as a similarity score over other metrics, because it deals best with data containing zero values. Second, we selected protein pairs that physically interact in living cells according to the BioGrid database (release version 4.4.209) (*26*). A total of 118 physical interactions were reported between proteins in our dataset. These interactions were detected with one of the following methods as reported by BioGrid database: Affinity Capture-MS, Affinity Capture-Western, Co-localization, Co-crystal structure, Co-purification, Co-fractionation, FRET, PCA, proximity label-MS and Two-hybrid. To determine whether the average cosine distance of the 118 interacting protein pairs was significantly different from that of random pairs of proteins, we compared it to the distribution of mean cosine distances obtained from 1000 randomly selected sets of 118 protein pairs.

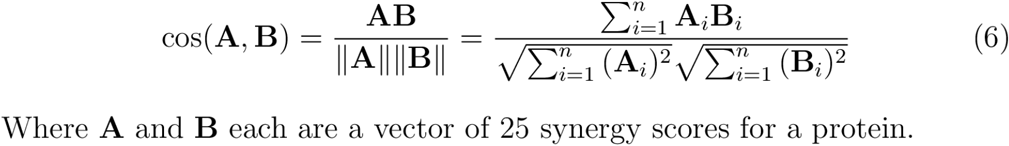

We also explored if proteins forming interaction cliques tend to have similar CCDs. To do so, we built an interaction network of physical interactions using the *igraph* package (version 1.3.4) and identified highest order cliques. We found three cliques with four elements each and displayed them on the UMAP plot (**Fig. 3C**).

#### Chromatin context dependent pathway activity in tumors

The hypothesis that we aimed to test, based on the observed CCD pattern of BRCA2 (**Fig. 2D**), is that loss of BRCA2 in human tumors should cause a shift of the MMEJ:NHEJ balance towards MMEJ specifically in a broad diversity of euchromatic regions and not in triple heterochromatin. For this we chose a recent whole-genome sequencing dataset derived from BRCA2-negative tumors from diverse tissue origins (n = 41) and HPV negative head and neck squamous cell carcinoma (HNSCC) samples (n = 22) (*41*). In the genomes of both tumor types we called small insertions and deletions (indels) and structural variants (SVs). We chose HPV-negative HNSCC as controls because they have a sufficiently high rate of indels and SVs to provide the required statistical power.

Indels were obtained from the final pan-cancer analysis of whole genomes (PCAWG) consortium somatic mutation list for the PCAWG-HNSCC (BRCA2^+/+^) and PCAWG-BRCA2^mut^ (BRCA2^-/-^) cohorts. The methods and post-calling filtering strategies were previously described in detail (*49*). Indels were subsequently classified using *indelsClassification* (https://github.com/ferrannadeu/indelsClassification) to identify deletions generated by error-prone NHEJ (>5 bp deletions without micro-homology), MMEJ repair (>5 bp deletions with ≥2 bp micro-homology sequence), polymerase slippage (1 bp deletions in ≥3 bp homopolymers) and indels in repeats (>1 bp deletions at ≥3 bp repeats) (**Table S9)**.

Additionally, we called structural variants (SVs) with BRASS (*50*) and annotated by AnnotateBRASS (https://github.com/MathijsSanders/AnnotateBRASS). We determined the following statistics per SV: the number of supporting read-pairs, the alignment position variance of supporting read-pairs, the frequency of read clipping, the frequency of reads with an excess of variants (≥ 2) absent from dbSNP, the proportion of read-pairs correctly oriented based on the SV detection and the number of SV-supporting read-pairs proximal to the SV breakpoints with alternative alignments (high genome homology). The post-annotation filtering strategy was previously described in detail (https://github.com/cancerit/BRASS). We analyzed the PCAWG-HNSCC (BRCA2^+/+^) and PCAWG-BRCA2^mut^ (BRCA2^-/-^) utilizing the same methodology (**Table S10**).

Next, we counted the total number of NHEJ or MMEJ small deletions in cLADs and ciLADs and calculated the log_2_(cLAD/ciLAD) ratio in each cohort. This ratio is a metric for the chromatin bias in the accumulation of NHEJ or MMEJ mutations in the different cohorts. We tested if the total number of NHEJ and MMEJ deletions were equally distributed between cLADs and ciLADs in each cohort with a two-sided Fisher’s exact test.

Additionally, we counted the total number of long MH deletions (size range 1.4 kb - 272.9 kb, 95% interval) contained within either a cLAD or ciLAD. We tested if the number of MH deletions were differently distributed between cLADs and ciLADs between cohorts by a two-sided Fisher’s exact test.

## SUPPLEMENTARY TABLES

**Table S1: KO gRNA library gRNA sequence list**.

Separate Excel file (AGM20191020_Table_1_DDR_library_IDT_SO#3121893.xlsx)

**Table S2:**
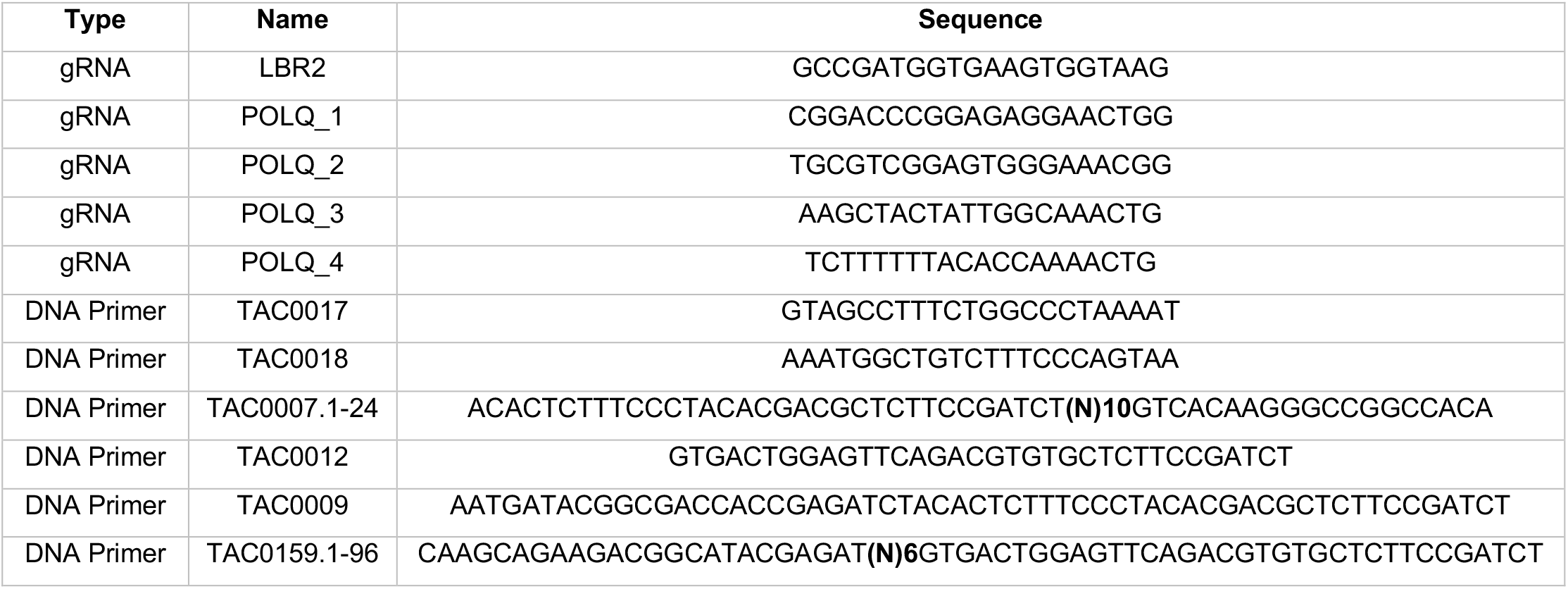
gRNA and primer sequences used in this manuscript.

**Table S4: Δlog**_**2**_**MMEJ:NHEJ scores for 519 proteins and 19 IPRs**.

Separate excel file (xv20220819_Table_S4_delta_log2_MMEJ_NHEJ.xlsx)

**Table S5:**
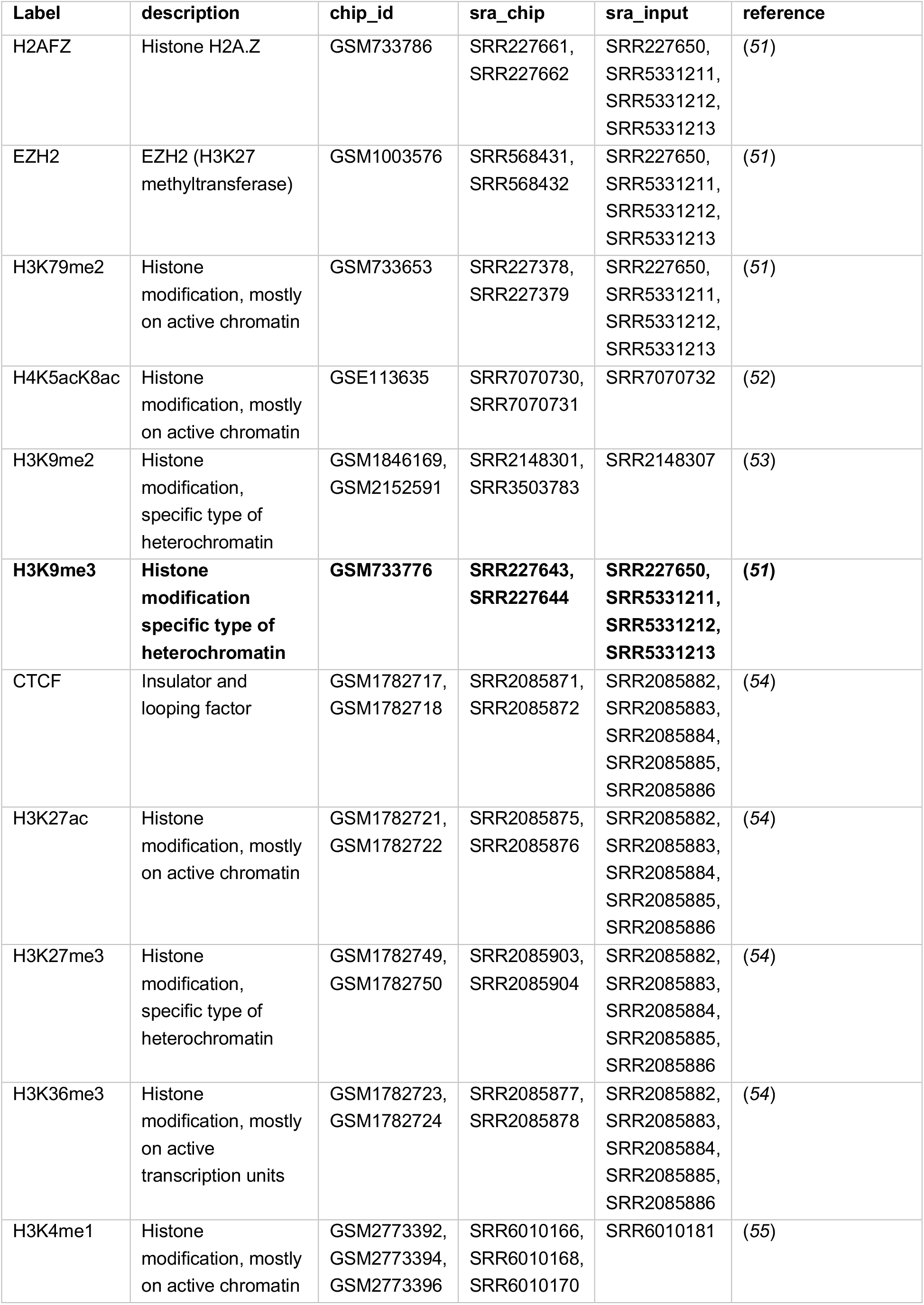
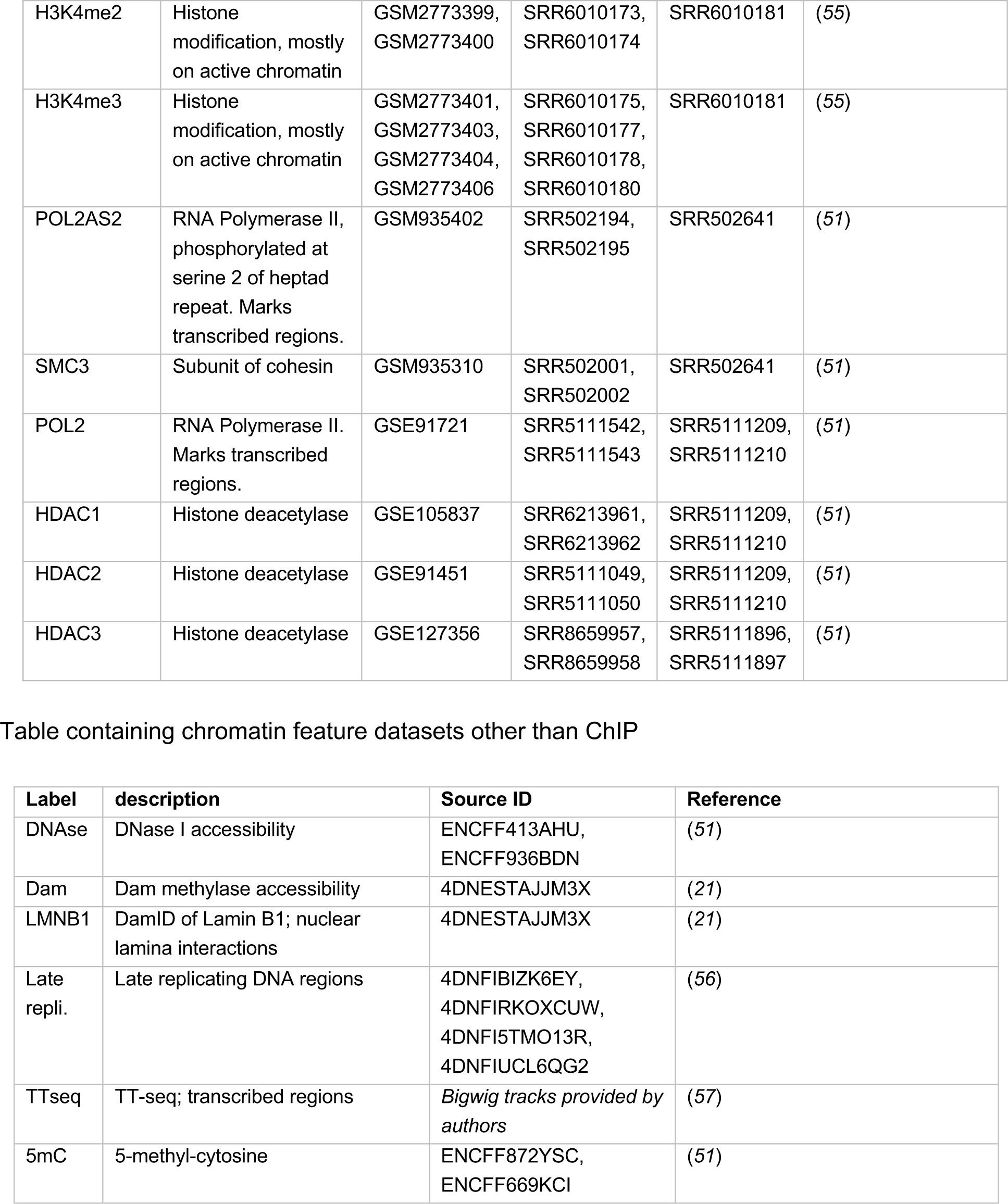
Epigenome maps. Data table adapted from (*6*). These are the epigenome Chromatin Immunoprecipitation (ChIP) datasets used in this study.

**Table S6: Genomic coordinates, chromatin feature scores and barcodes of 19 IPRs in K562 clone 5**

Separate excel file (xv20220819_Table_S6_clone_5_chromatin_features.xlsx)

**Table S7: Chromatin context dependent effects of proteins in the screen**.

Separate excel file (xv20220929_Table_S7_global_CCD_MMEJ_NHEJ_results.xlsx)

**Table S8: CCD similarity scores of proteins with physical interactions (curated by BioGrid)**

Separate excel file (xv20220913_Table_S8_BioGRID_interaction_CCD.xlsx)

**Table S9: Mapped indels in BRCA2**^**+/+**^ **and BRCA2**^**-/-**^ **tumors**.

Separate excel file (xv20220922_Table_S9_indel_mutations_tumors_COSMIC.xlsx)

**Table S10: Mapped structural variants in BRCA2**^**+/+**^ **and BRCA2**^**-/-**^ **tumors**.

Separate excel file (xv20220922_Table_S10_SV_mutations_tumors_BRASS.xlsx)

## SUPPLEMENTARY FIGURES

**Fig. S1:**
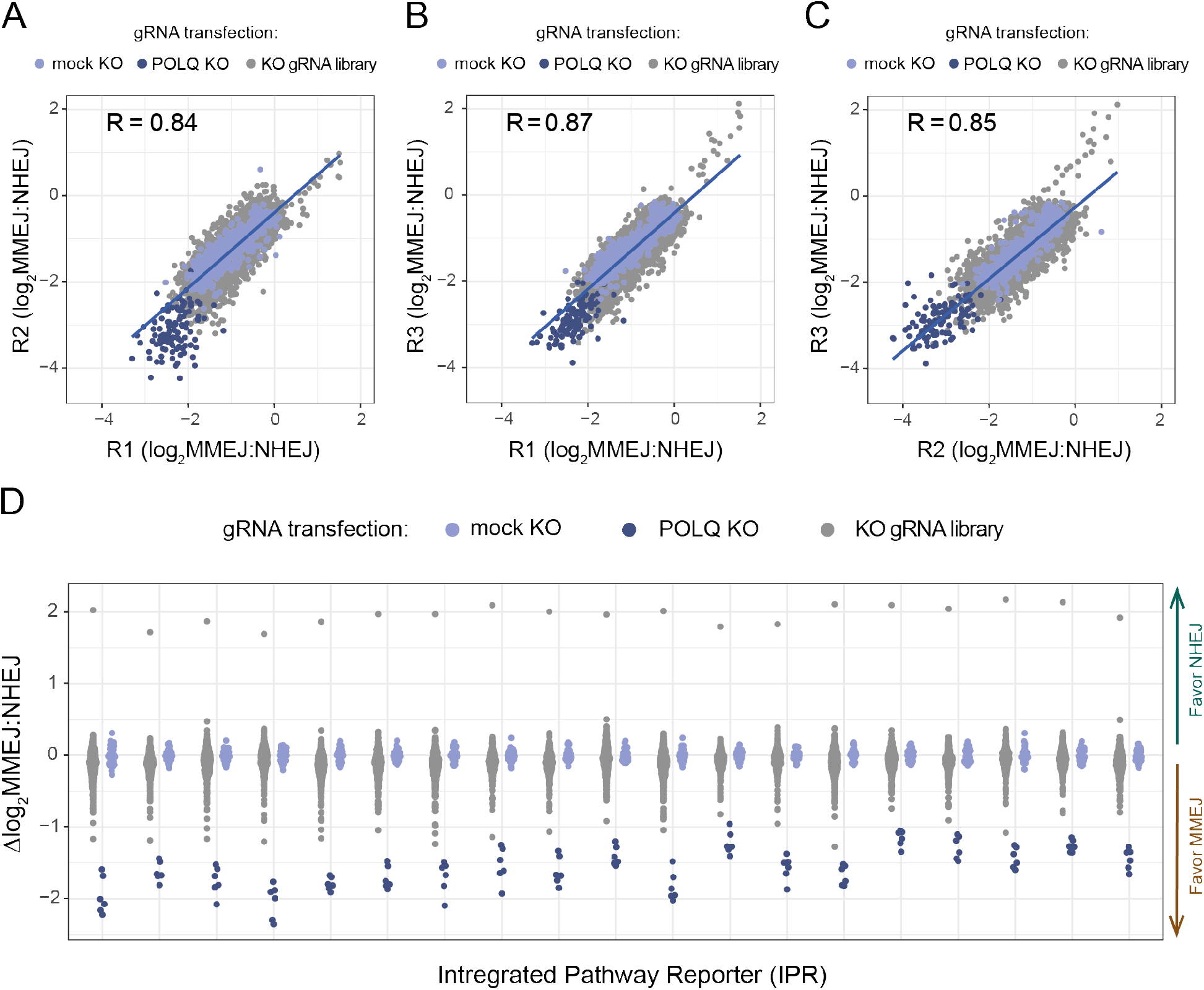
Screen replicate reproducibility and distribution of Δlog_2_MMEJ:NHEJ values. A-C) Pairwise correlations of log_2_MMEJ:NHEJ values of individual IPRs between replicate experiments R1, R2 and R3, after application of quality filters as described in step 3 of the data processing. R denotes Pearson correlation coefficient. D) Dynamic range of Δlog_2_MMEJ:NHEJ balances after averaging of replicates.

**Fig. S2:**
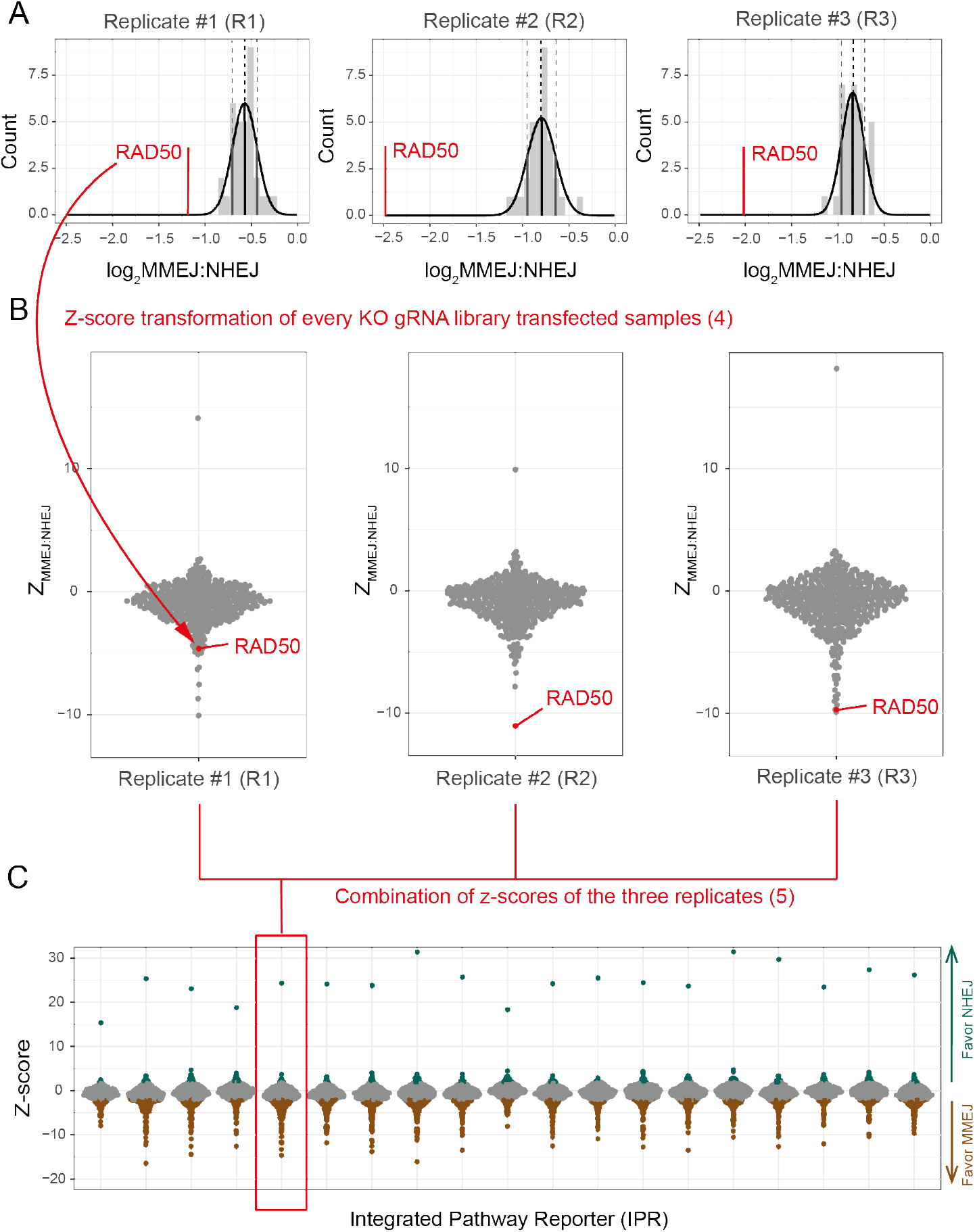
z-transformation and combining of replicate measurements of log_2_MMEJ:NHEJ values. A) Histogram of log_2_MMEJ:NHEJ balance of mock KO transfected samples of a single IPR (IPR_barcode: CATTTCTGATCAATAA). The fitted normal distribution is depicted in black. Mean (black) and mean ± one standard deviation (grey) highlighted with vertical dotted lines. In red, log_2_MMEJ:NHEJ balance of RAD50 KO is plotted as an example to illustrate the z-score transformation for a single protein. A red arrow is displayed connecting RAD50 KO data point in replicate #1 panel A and B. Each panel represents a different replicate and a similar arrow could be drawn for the other replicates as well. B) Beeswarm plot of the z-score transformed log_2_MMEJ:NHEJ balance of KO samples for a single reporter (CATTTCTGATCAATAA) (Formula 4). C) z-score transformed log_2_MMEJ:NHEJ balance perturbations after combining three replicates for every MMEJ:NHEJ pathway reporters by the Stouffer’s method (Formula 5). A value outside the [-1.96,1.96] range is considered to be significant with a significance level of >95%. Positive values represent proteins that *favor NHEJ* (green dots and arrow) and negative values proteins that *favor MMEJ* (brown dots and arrow).

**Fig. S3:**
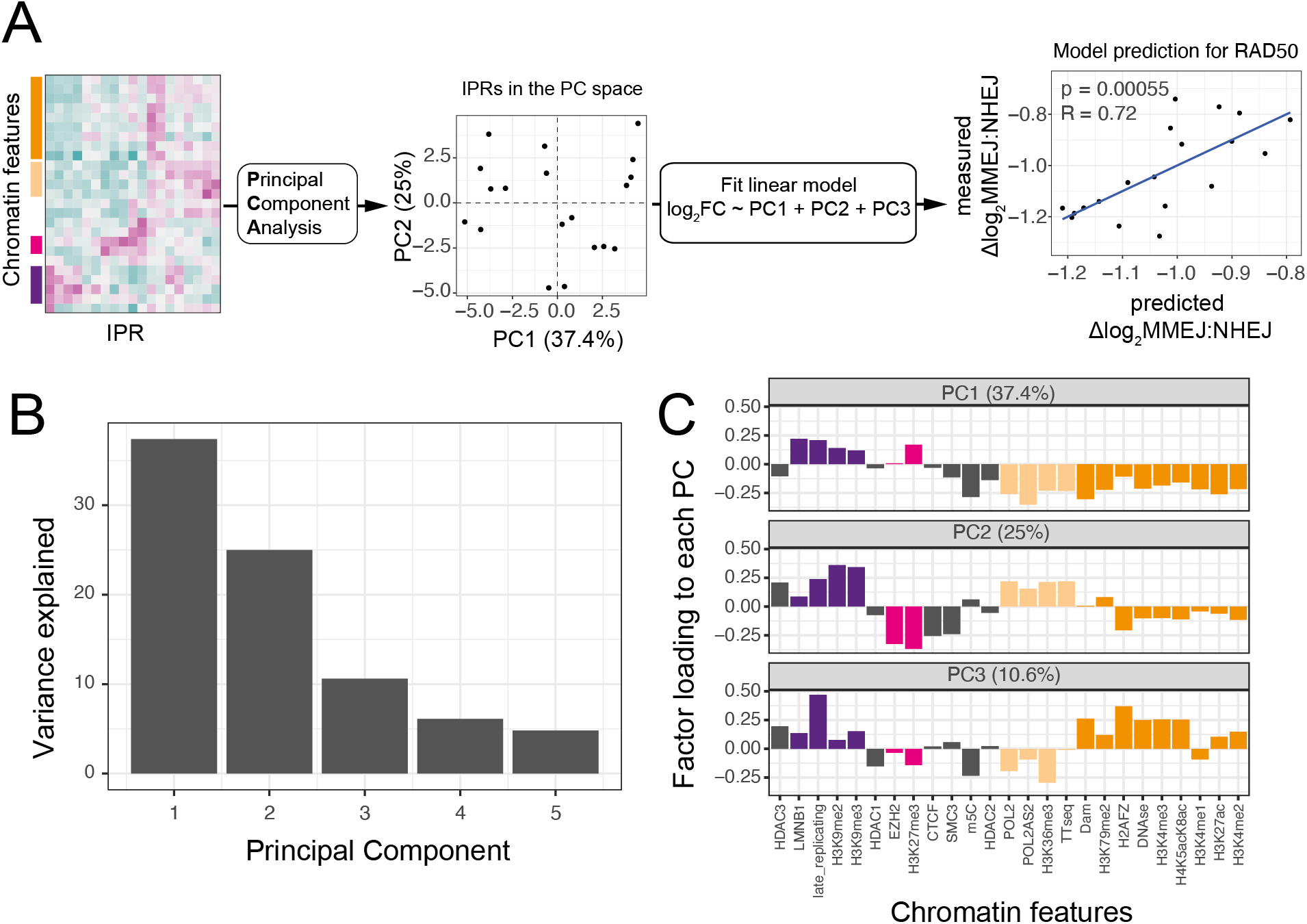
Principal component regression analysis. A) Principal Component Regression workflow. First, a principal component analysis (PCA) was run with 25 chromatin feature values for each IPR. Second, exploration of the PCA revealed that the first three PCs recapitulate most of the chromatin feature variance. Third, a linear model with three principal components was ran for each protein and the performance was assessed by predicted vs. measured comparison. B) Percentage of variance explained by the first five PCs. C) Bar graph showing the weight of each chromatin feature for PC1, PC2 and PC3. Bars are coloured according to the chromatin context they represent: Triple heterochromatin (purple), H3K27me3 heterochromatin (pink), transcription (light orange) and enhancers/promoters (orange).

**Fig. S4:**
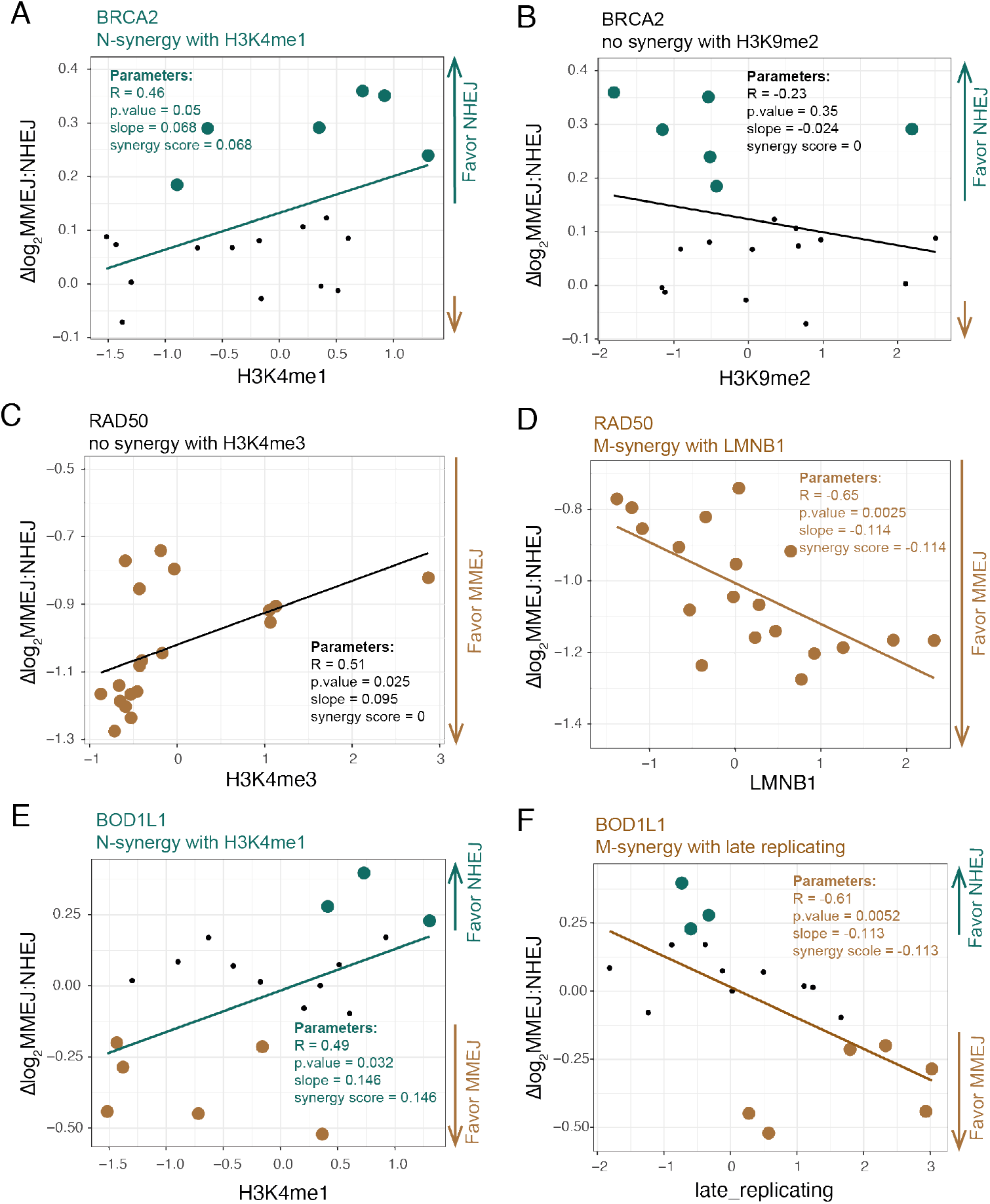
Examples of linear fit correlation with individual chromatin features. Examples of three proteins with significant CCDs on individual chromatin features. All correlation plots show data of the 19 IPRs, the linear regression fit, and regression analysis parameters that are relevant for the *synergy score*. (R = Pearson correlation coefficient, p.value = p value of the correlation coefficient, slope = slope of the linear fit, synergy score = final value of CCD interaction between protein and chromatin feature after corrections). Color scheme of the figure shows if the protein - chromatin feature has an M-synergy (brown), N-synergy (green) or no synergy (black). A) N-synergy between BRCA2 and H3K4me1. B) No synergy between BRCA2 and H3K9me2. This interaction is explained by the absence of the chromatin feature and therefore is discarded (*favor NHEJ* and slope < 0). C) No synergy between RAD50 and H3K4me3. Same as for B applies here (*favor MMEJ* and slope > 0). D) M-synergy between RAD50 and interactions with the nuclear lamina (LMNB1). E) N-synergy between BOD1L and H3K4me1. F) M-synergy between BOD1L and late replicating chromatin.

**Fig. S5:**
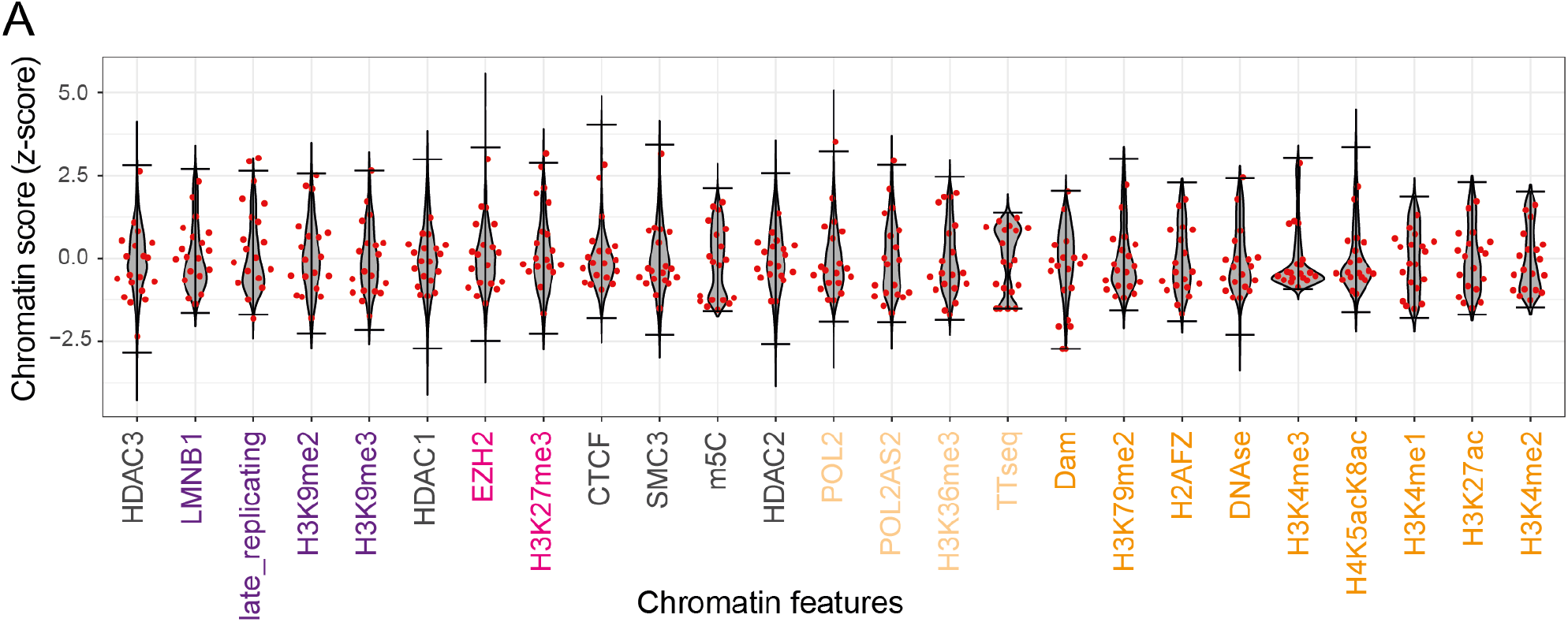
Estimation of genome-wide dynamic ranges of chromatin features. In grey, distribution of genome-wide chromatin scores, with the 99% confidence interval (99CI) marked by the horizontal black lines (top 0.5% and bottom 0.5%). In red, chromatin scores of all 19 IPRs in clone 5 for which MMEJ:NHEJ balance was measured in the screen.

**Fig. S6:**
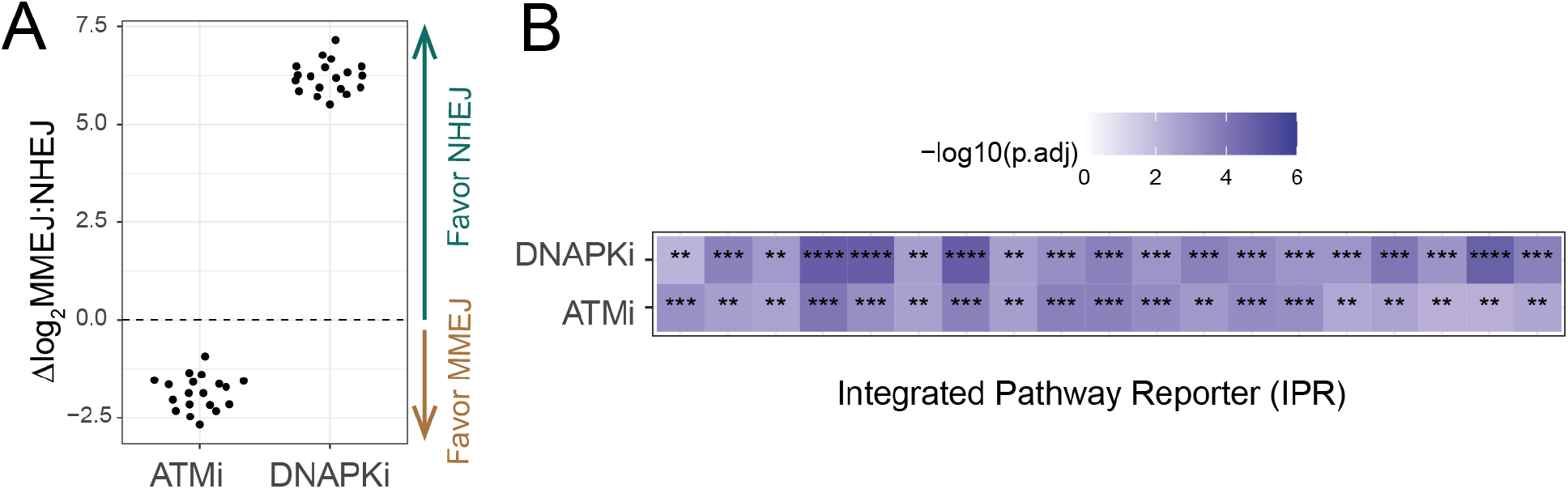
ATM and DNAPK inhibitor effects. A) Δlog_2_MMEJ:NHEJ of ATM and DNAPKcs inhibitor of each IPR. B) Adjusted p-values of Student’s t-test comparing Δlog_2_MMEJ:NHEJ scores in ATM and DNAPKcs inhibited compared to the vehicle control (n = 3) for each reporter. This test was used to confirm significance of the changes in the log_2_MMEJ:NHEJ.

**Fig. S7:**
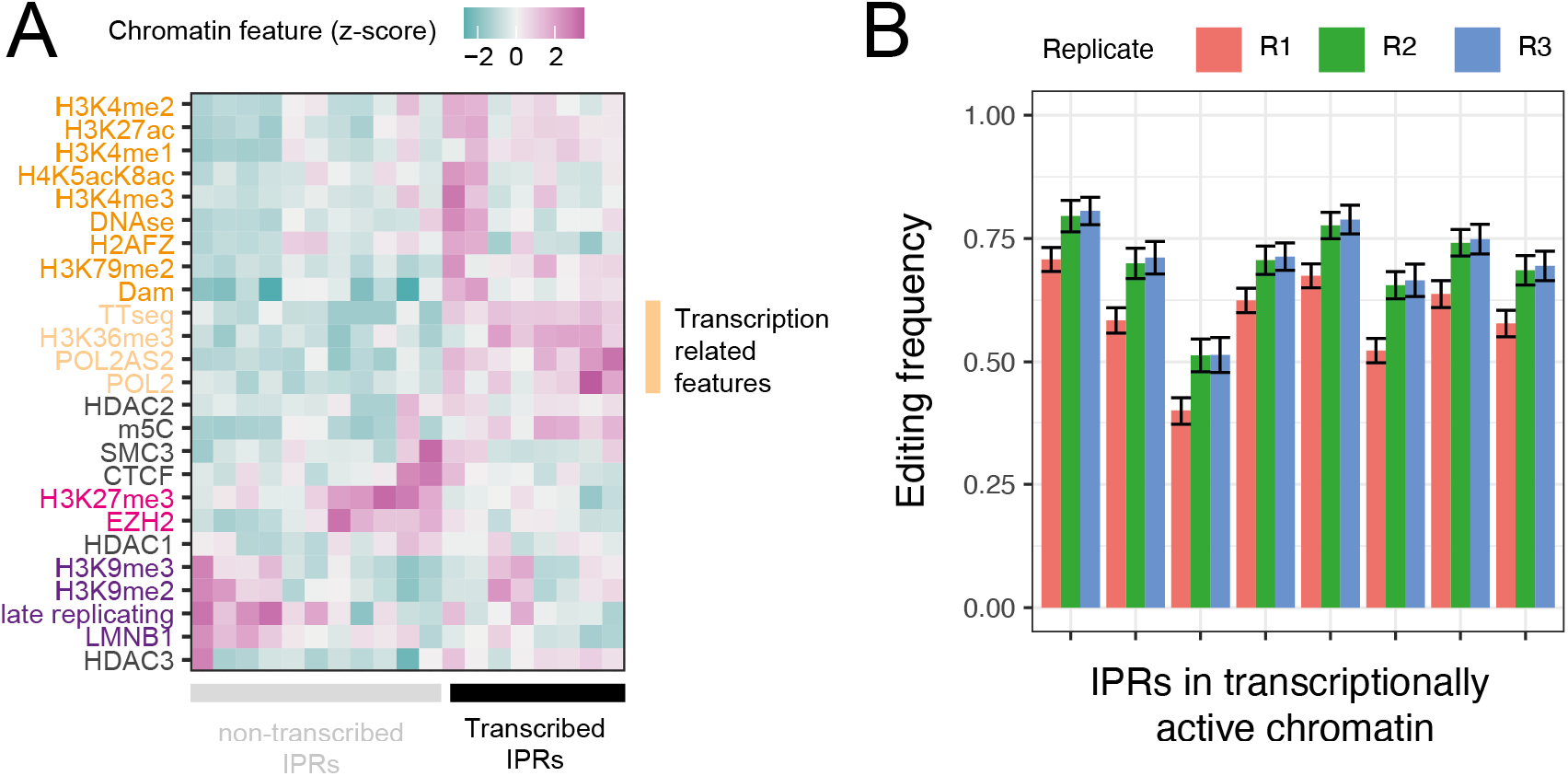
Editing efficiency estimation. A) Transcribed (n = 8) and non-transcribed (n = 11) IPRs in clone 5. We classified IPRs based on the transcription-related feature signals (light orange). B) Editing efficiencies in transcribed IPRs for each replicate in mock transfected samples (n = 33). Error bars show mean ± sd.

**Fig. S8:**
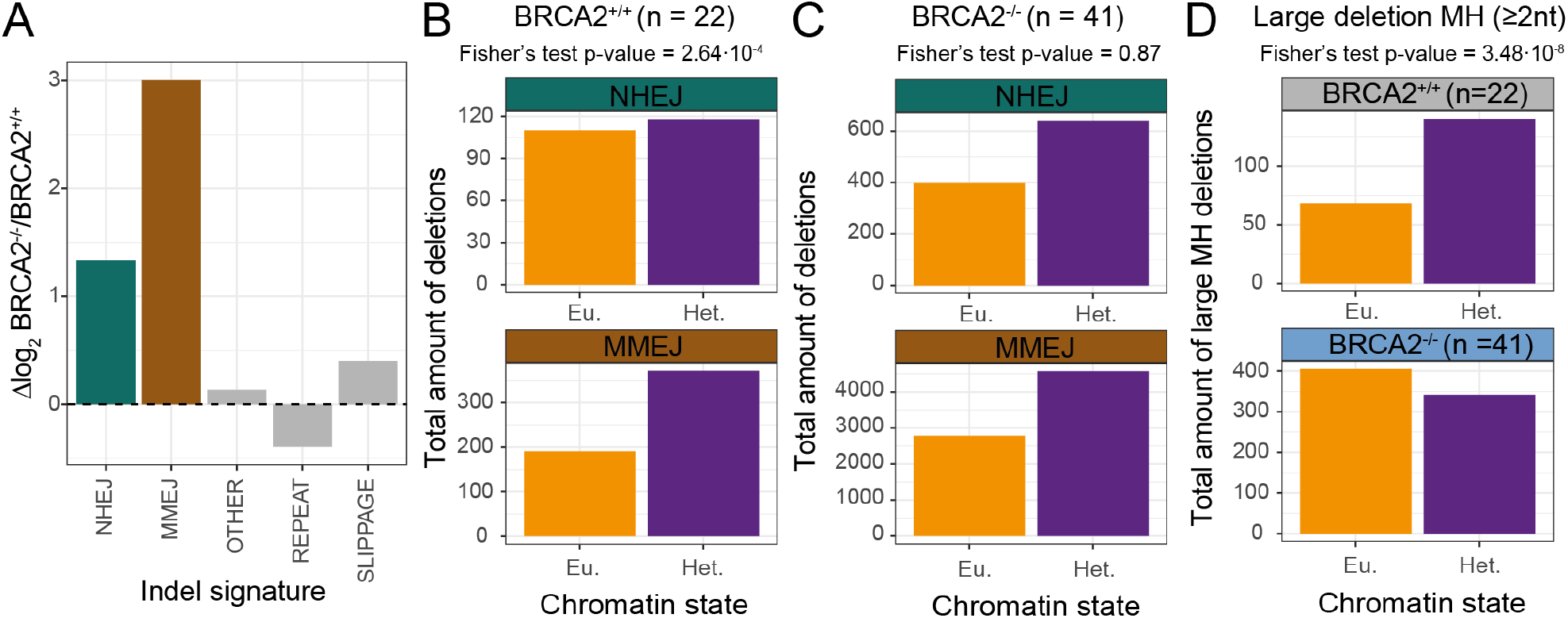
BRCA2^+/+^ and BRCA2^-/-^ mutation analysis. (**A**) log_2_ fold change of average frequencies of the five short indel signatures (see Methods) called per tumor in BRCA2^-/-^ compared to BRCA2^+/+^. (**B-C**) Distribution of NHEJ and MMEJ counts over euchromatin (Eu.) and constitutive lamina-associated heterochromatin (Het.) in BRCA2^+/+^ tumors (B) and BRCA2^-/-^ tumors (C). For each tumor type a Fisher’s exact test was applied to test for differential distribution of NHEJ and MMEJ counts between Het and Eu.(**D**) Similar analysis as in B-C bottom rows, showing total number of large deletions with MH at break sites in euchromatin (Eu.) and constitutive lamina-associated heterochromatin (Het.) in BRCA2^+/+^ and BRCA2^-/-^ tumors.

## REFERENCES

1. R. Scully, A. Panday, R. Elango, N. A. Willis, DNA double-strand break repair-pathway choice in somatic mammalian cells. Nat Rev Mol Cell Biol 20, 698–714 (2019).

2. N. J. O’Neil, M. L. Bailey, P. Hieter, Synthetic lethality and cancer. Nat Rev Genet 18, 613–623 (2017).

3. N. Hustedt, D. Durocher, The control of DNA repair by the cell cycle. Nat Cell Biol 19, 1–9 (2016).

4. A. Schipler, G. Iliakis, DNA double-strand-break complexity levels and their possible contributions to the probability for error-prone processing and repair pathway choice. Nucleic Acids Res 41, 7589–7605 (2013).

5. F. Aymard et al., Transcriptionally active chromatin recruits homologous recombination at DNA double-strand breaks. Nat Struct Mol Biol 21, 366–374 (2014).

6. R. Schep et al., Impact of chromatin context on Cas9-induced DNA double-strand break repair pathway balance. Mol Cell 81, 2216–2230 e2210 (2021).

7. S. L. Sanders et al., Methylation of histone H4 lysine 20 controls recruitment of Crb2 to sites of DNA damage. Cell 119, 603–614 (2004).

8. Y. Huyen et al., Methylated lysine 79 of histone H3 targets 53BP1 to DNA double-strand breaks. Nature 432, 406–411 (2004).

9. Y. Sun et al., Histone H3 methylation links DNA damage detection to activation of the tumour suppressor Tip60. Nat Cell Biol 11, 1376–1382 (2009).

10. M. van Overbeek et al., DNA Repair Profiling Reveals Nonrandom Outcomes at Cas9-Mediated Breaks. Mol Cell 63, 633–646 (2016).

11. D. Setiaputra, D. Durocher, Shieldin - the protector of DNA ends. EMBO Rep 20, (2019).

12. Y. Xie et al., RBX1 prompts degradation of EXO1 to limit the homologous recombination pathway of DNA double-strand break repair in G1 phase. Cell Death Differ 27, 1383–1397 (2020).

13. F. Robert, M. Barbeau, S. Ethier, J. Dostie, J. Pelletier, Pharmacological inhibition of DNA-PK stimulates Cas9-mediated genome editing. Genome Med 7, 93 (2015).

14. V. T. Chu et al., Increasing the efficiency of homology-directed repair for CRISPR-Cas9-induced precise gene editing in mammalian cells. Nat Biotechnol 33, 543–548 (2015).

15. C. J. Tsai, S. A. Kim, G. Chu, Cernunnos/XLF promotes the ligation of mismatched and noncohesive DNA ends. Proc Natl Acad Sci U S A 104, 7851–7856 (2007).

16. A. Craxton et al., PAXX and its paralogs synergistically direct DNA polymerase lambda activity in DNA repair. Nat Commun 9, 3877 (2018).

17. J. A. Hussmann et al., Mapping the genetic landscape of DNA double-strand break repair. Cell 184, 5653–5669 e5625 (2021).

18. S. M. Howard, D. A. Yanez, J. M. Stark, DNA damage response factors from diverse pathways, including DNA crosslink repair, mediate alternative end joining. PLoS Genet 11, e1004943 (2015).

19. W. Akhtar et al., Chromatin position effects assayed by thousands of reporters integrated in parallel. Cell 154, 914–927 (2013).

20. M. Corrales et al., Clustering of Drosophila housekeeping promoters facilitates their expression. Genome Res 27, 1153–1161 (2017).

21. C. Leemans et al., Promoter-Intrinsic and Local Chromatin Features Determine Gene Repression in LADs. Cell 177, 852–864 e814 (2019).

22. A. Sfeir, L. S. Symington, Microhomology-Mediated End Joining: A Back-up Survival Mechanism or Dedicated Pathway? Trends Biochem Sci 40, 701–714 (2015).

23. P. Khongkow et al., FOXM1 targets NBS1 to regulate DNA damage-induced senescence and epirubicin resistance. Oncogene 33, 4144–4155 (2014).

24. M. Kriegs et al., The epidermal growth factor receptor modulates DNA double-strand break repair by regulating non-homologous end-joining. DNA Repair (Amst) 9, 889–897 (2010).

25. L. Wan et al., Scaffolding protein SPIDR/KIAA0146 connects the Bloom syndrome helicase with homologous recombination repair. Proc Natl Acad Sci U S A 110, 10646–10651 (2013).

26. C. Stark et al., BioGRID: a general repository for interaction datasets. Nucleic Acids Res 34, D535–539 (2006).

27. A. A. Goodarzi et al., ATM signaling facilitates repair of DNA double-strand breaks associated with heterochromatin. Mol Cell 31, 167–177 (2008).

28. I. Chiolo et al., Double-strand breaks in heterochromatin move outside of a dynamic HP1a domain to complete recombinational repair. Cell 144, 732–744 (2011).

29. S. Matsuoka et al., Ataxia telangiectasia-mutated phosphorylates Chk2 in vivo and in vitro. Proc Natl Acad Sci U S A 97, 10389–10394 (2000).

30. Y. Shiloh, Y. Ziv, The ATM protein kinase: regulating the cellular response to genotoxic stress, and more. Nat Rev Mol Cell Biol 14, 197–210 (2013).

31. S. Choi, A. M. Gamper, J. S. White, C. J. Bakkenist, Inhibition of ATM kinase activity does not phenocopy ATM protein disruption: implications for the clinical utility of ATM kinase inhibitors. Cell Cycle 9, 4052–4057 (2010).

32. G. P. Crossan, K. J. Patel, The Fanconi anaemia pathway orchestrates incisions at sites of crosslinked DNA. J Pathol 226, 326–337 (2012).

33. J. Torres-Rosell et al., The Smc5-Smc6 complex and SUMO modification of Rad52 regulates recombinational repair at the ribosomal gene locus. Nat Cell Biol 9, 923–931 (2007).

34. R. Bayley et al., H3K4 methylation by SETD1A/BOD1L facilitates RIF1-dependent NHEJ. Mol Cell 82, 1924–1939 e1910 (2022).

35. M. Hoek, M. P. Myers, B. Stillman, An analysis of CAF-1-interacting proteins reveals dynamic and direct interactions with the KU complex and 14-3-3 proteins. J Biol Chem 286, 10876–10887 (2011).

36. T. S. Nambiar, L. Baudrier, P. Billon, A. Ciccia, CRISPR-based genome editing through the lens of DNA repair. Mol Cell 82, 348–388 (2022).

37. L. Feng, J. Wang, J. Chen, The Lys63-specific deubiquitinating enzyme BRCC36 is regulated by two scaffold proteins localizing in different subcellular compartments. J Biol Chem 285, 30982–30988 (2010).

38. Y. Hu et al., RAP80-directed tuning of BRCA1 homologous recombination function at ionizing radiation-induced nuclear foci. Genes Dev 25, 685–700 (2011).

39. S. Ahrabi et al., A role for human homologous recombination factors in suppressing microhomology-mediated end joining. Nucleic Acids Res 44, 5743–5757 (2016).

40. J. Zamborszky et al., Loss of BRCA1 or BRCA2 markedly increases the rate of base substitution mutagenesis and has distinct effects on genomic deletions. Oncogene 36, 746–755 (2017).

41. A. L. H. Webster et al., Fanconi Anemia Pathway Deficiency Drives Copy Number Variation in Squamous Cell Carcinomas. bioRxiv, 2021.2008.2014.456365 (2021).

42. T. Clouaire, G. Legube, A Snapshot on the Cis Chromatin Response to DNA Double-Strand Breaks. Trends Genet 35, 330–345 (2019).

43. E. K. Brinkman et al., Kinetics and Fidelity of the Repair of Cas9-Induced Double-Strand DNA Breaks. Mol Cell 70, 801–813 e806 (2018).

44. E. K. Brinkman, T. Chen, M. Amendola, B. van Steensel, Easy quantitative assessment of genome editing by sequence trace decomposition. Nucleic Acids Res 42, e168 (2014).

45. A. Hendel et al., Quantifying genome-editing outcomes at endogenous loci with SMRT sequencing. Cell Rep 7, 293–305 (2014).

46. R. Schep, C. Leemans, E. K. Brinkman, T. van Schaik, B. van Steensel, Protocol: A Multiplexed Reporter Assay to Study Effects of Chromatin Context on DNA Double-Strand Break Repair. Front Genet 12, 785947 (2021).

47. A. M. Yu, M. McVey, Synthesis-dependent microhomology-mediated end joining accounts for multiple types of repair junctions. Nucleic Acids Res 38, 5706–5717 (2010).

48. S. H. Chan, A. M. Yu, M. McVey, Dual roles for DNA polymerase theta in alternative end-joining repair of double-strand breaks in Drosophila. PLoS Genet 6, e1001005 (2010).

49. L. Moore et al., The mutational landscape of normal human endometrial epithelium. Nature 580, 640–646 (2020).

50. Icgc Tcga Pan-Cancer Analysis of Whole Genomes Consortium, Pan-cancer analysis of whole genomes. Nature 578, 82–93 (2020).

51. Encode Project Consortium, An integrated encyclopedia of DNA elements in the human genome. Nature 489, 57–74 (2012).

52. C. J. Ott et al., Enhancer Architecture and Essential Core Regulatory Circuitry of Chronic Lymphocytic Leukemia. Cancer Cell 34, 982–995 e987 (2018).

53. A. C. Salzberg et al., Genome-wide mapping of histone H3K9me2 in acute myeloid leukemia reveals large chromosomal domains associated with massive gene silencing and sites of genome instability. PLoS One 12, e0173723 (2017).

54. C. Schmidl, A. F. Rendeiro, N. C. Sheffield, C. Bock, ChIPmentation: fast, robust, low-input ChIP-seq for histones and transcription factors. Nat Methods 12, 963–965 (2015).

55. R. N. Shah et al., Examining the Roles of H3K4 Methylation States with Systematically Characterized Antibodies. Mol Cell 72, 162–177 e167 (2018).

56. J. Dekker et al., The 4D nucleome project. Nature 549, 219–226 (2017).

57. B. Schwalb et al., TT-seq maps the human transient transcriptome. Science 352, 1225–1228 (2016).

